# PUF family proteins FBF-1 and FBF-2 regulate germline stem and progenitor cell proliferation and differentiation in *C. elegans*

**DOI:** 10.1101/825984

**Authors:** Xiaobo Wang, Mary Ellenbecker, Benjamin Hickey, Nicholas J. Day, Ekaterina Voronina

## Abstract

Stem cells support tissue maintenance, but the mechanisms that balance the rate of stem cell self-renewal with differentiation at a population level remain uncharacterized. Through investigating the regulation of germline stem cells by two PUF family RNA-binding proteins FBF-1 and FBF-2 in *C. elegans*, we find that FBF-1 restricts differentiation, while FBF-2 promotes both proliferation and differentiation. FBFs act on a shared set of target mRNAs; however, FBF-1 destabilizes target transcripts, while FBF-2 promotes their accumulation. These regulatory differences result in complementary effects of FBFs on stem cells. We identify a mitotic cyclin as one of the targets affecting stem cell homeostasis. FBF-1-mediated translational control requires the activity of CCR4-NOT deadenylase. Distinct abilities of FBFs to cooperate with CCR4-NOT depend on protein sequences outside of the conserved PUF family RNA-binding domain. We propose that the combination of FBF activities regulates the dynamics of germline stem cell proliferation and differentiation.

## INTRODUCTION

Adult tissue maintenance relies on the activity of stem cells that self-renew and produce differentiating progeny in step with tissue demands (Morrison and Kimble 2006). It is essential that self-renewal be balanced with differentiation to preserve the size of the stem cell pool over time. One simple model achieving this balance is an asymmetric division that always produces a single stem cell daughter and a daughter destined to differentiate (Chen and others 2016). Alternatively, tissue homeostasis can be controlled at a population level (Simons and Clevers 2011), where some stem cells are lost through differentiation while others proliferate, with both outcomes occurring with the same frequency. Such population-level control of stem cell activity is observed in the *C. elegans* germline (Kimble and Crittenden 2007). However, the mechanisms of population-level balance of stem cell proliferation and differentiation in the adult tissues are largely unclear.

The *C. elegans* hermaphrodite germline is a robust system to explore the mechanisms coordinating stem cell proliferation and differentiation. It is maintained by a stem cell niche that supports about 200-250 mitotically-dividing <u>s</u>tem and <u>p</u>rogenitor <u>c</u>ells at the distal end of the gonad (collectively called SPCs, **Figure 1A, Cii**). A single somatic distal tip cell serves as a stem cell niche and activates the GLP-1/Notch signaling necessary for SPC pool maintenance (Austin and Kimble 1987), which in turn supports germline development (Hansen and Schedl 2013). As germline stem cells move proximally away from the niche, they differentiate by entering meiotic prophase and eventually generate gametes near the proximal gonad end. Mitotic divisions of SPCs are not oriented and there doesn’t appear to be a correlation between the position of cell divisions uniformly distributed over the SPC zone and the position of cells committing to differentiation at the proximal end of the zone (Crittenden and others 2006).

**Figure 1.**
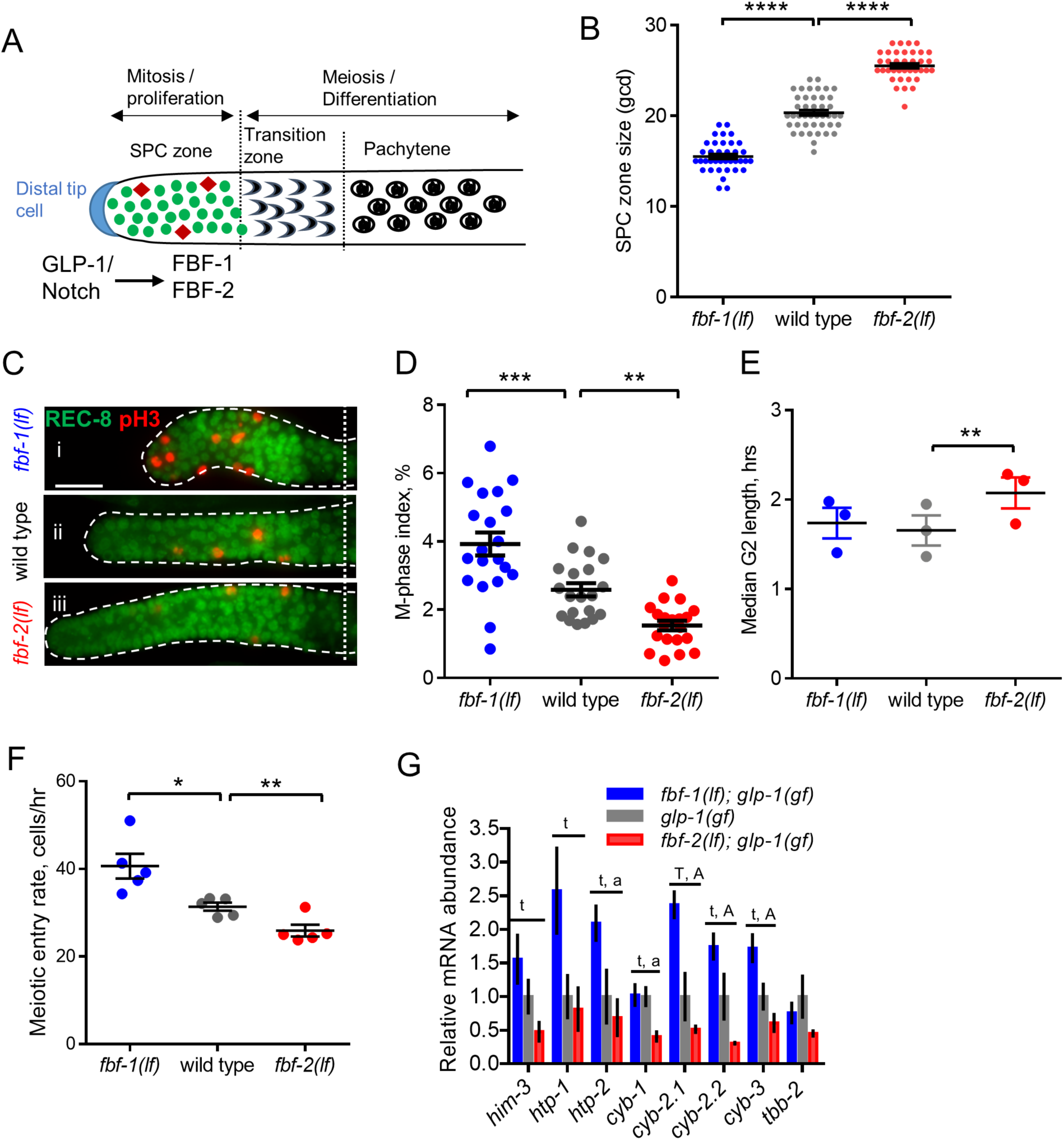
FBF-1 and FBF-2 differentially regulate germline <u>s</u>tem and <u>p</u>rogenitor <u>c</u>ell (SPC) zone size. (A) Schematic of the distal germline of *C. elegans* adult hermaphrodite. GLP-1/Notch signaling from the distal tip cell (blue) supports germline SPC proliferation. Progenitors enter meiosis when they reach the transition zone. FBF-1 and FBF-2, downstream of GLP-1/Notch, are required for SPC maintenance. Green circles, stem and progenitor cells; red diamonds, mitotically dividing cells. (B) SPC zone sizes of the wild type, *fbf-1(lf)* and *fbf-2(lf)* germlines were measured by counting germ cell diameters (gcd) spanning SPC zone. Genetic background is indicated on the X-axis and SPC zone size on the Y-axis. (C) Distal germlines dissected from adult wild type, *fbf-1(lf)*, and *fbf-2(lf)* hermaphrodites and stained with anti-REC-8 (green) and anti-<u>p</u>hospho-Histone <u>H3</u> (pH3; red) to visualize the SPC zone and mitotic cells in M-phase. Germlines are outlined with the dashed lines and the vertical dotted line marks the beginning of transition zone as recognized by the ‘crescent-shaped’ chromatin and loss of REC-8. Scale bar: 10 μm. (D) Quantification of mitotic indices of germline SPCs in animals of different genetic backgrounds (as indicated on the X-axis). (B, D) Plotted values are individual data points and arithmetical means ± S.E.M. Differences in SPC pool sizes and mitotic indices were evaluated by one-way ANOVA with Dunnett’s post-test. Data were collected from 3 independent experiments and 20-43 germlines were scored for each genotype. (E) Median SPC G2-phase length in different genetic backgrounds (as indicated on the X-axis). Plotted values are individual data points and arithmetical means ± S.E.M. Difference in median G2 length was evaluated by one-way ANOVA with Dunnett’s post-test. Data were collected from 3 independent experiments as shown in Figure 1—figure supplement 1B. (F) Meiotic entry rate of germline progenitors in different genetic backgrounds (as indicated on the X-axis). Plotted values are individual data points and arithmetical means ± S.E.M. Differences in meiotic entry rate between each *fbf* and the wild type were evaluated by one-way ANOVA with T-test with Bonferroni correction post-test. Data were collected from 5 independent experiments as shown in Figure 1—figure supplement 1C. (B-F) Asterisks mark statistically-significant differences (****, *P*<0.0001; ***, *P*<0.001; **, *P*<0.01; *, *P*<0.05). (G) Steady-state mRNA abundance of FBF target genes and a control non-FBF target gene in *glp-1(gf), glp-1(gf); fbf-1(lf)* and *glp-1(gf); fbf-2(lf)* genetic backgrounds was determined by qRT-PCR and normalized to the levels of actin (*act-1*). Tested FBF target genes are associated with meiotic entry (*htp-1, htp-2* and *him-3*) or cell cycle regulation (*cyb-1, cyb-2.1, cyb-2.2* and *cyb-3*). The control gene is a tubulin subunit, *tbb-2*. Reported abundance represents the arithmetical mean ± S.E.M of 3 independent biological replicates. Differences in mRNA abundance were evaluated by one-way ANOVA tests with linear trend and Tukey’s post-tests. Test for linear trend between column means (left to right): t, *P* <0.05; T, *P*<0.01. Tukey’s test for column means difference: a, *P*<0.05; A, *P* <0.01.

Analysis of *C. elegans* germline stem cell maintenance identified a number of genes affecting SPC self-renewal and differentiation (Hansen and Schedl 2013). Genes essential for self-renewal include GLP-1/Notch and two highly similar Pumilio and FBF (PUF) family RNA-binding proteins called FBF-1 and FBF-2 (Austin and Kimble 1987; Crittenden and others 2002; Zhang and others 1997). Genetic studies of stem cell maintenance led to a model where a balance of mitosis- and meiosis-promoting activities maintains tissue homeostasis (Hansen and Schedl 2013), but the regulatory mechanism matching differentiation demands with proliferative SPC activity remained elusive.

Importantly, SPC cell cycle is distinct from that of most somatic stem cells. One characteristic feature of *C. elegans* germline SPC cell cycle is a very short G1 phase (Fox and others 2011; Furuta and others 2018), reminiscent of the short G1 phase observed in embryonic stem cells (ESCs, (Becker and others 2006; Kareta and others 2015; White and Dalton 2005). Mouse and human ESCs maintain robust proliferation supported by cell cycle with a short G1 phase while the length of S and G2 phases is similar to that observed in differentiated mouse somatic cells (Becker and others 2006; Chao and others 2019; Kareta and others 2015; Stead and others 2002). Despite the abbreviated G1 phase, ESCs maintain S and G2 checkpoints (Chuykin and others 2008; Stead and others 2002; White and Dalton 2005). Similarly, *C. elegans* SPCs retain G2 checkpoints despite the shortened G1 phase (Garcia-Muse and Boulton 2005; Seidel and Kimble 2015). This may be due to a constant proliferative demand that both SPCs and ESCs are subject to. By contrast, this type of modified cell cycle is not observed in the adult stem cell populations that support regenerative response upon injury, such as adult mammalian bulge stem cells (hair follicle stem cells; (Cotsarelis and others 1990) or satellite cells (muscle stem cells; (Schultz 1974; 1985; Snow 1977) that remain in G0 or quiescent phase for the most of the adult life and only reenter cell cycle upon injury. Similarly, adult epidermal stem cells maintaining tissue homeostasis regulate their cell cycle by controlling G1/S transition (Mesa and others 2018).

Unlike somatic cells’ G1 phase that is triggered and marked by increased amounts of cyclins E and D (Aleem and others 2005; Guevara and others 1999), the germ cells characterized by a shortened G1 phase maintain a constitutive robust expression of G1/S regulators Cyclin E and CDK2 (Fox and others 2011; Furuta and others 2018; White and Dalton 2005). Despite continuous proliferation of *C. elegans* SPCs, the rate of SPC proliferation changes during development and in different mutant backgrounds (Michaelson and others 2010; Roy and others 2016) and a mechanism for changing the rate of proliferation to meet the demands of germ cell production while maintaining cell cycle with an abbreviated G1 phase remains unknown. Here, we report the mechanism through which PUF family RNA binding proteins FBF-1 and FBF-2 balance SPC proliferative activity with the rate of meiotic entry.

PUF proteins are expressed in germ cells of many animals and are conserved regulators of stem cells (Salvetti and others 2005; Wickens and others 2002). *C. elegans* PUF proteins expressed in germline SPCs, FBF-1 and FBF-2, share the majority of their target mRNAs (Porter and others 2019; Prasad and others 2016) and are redundantly required for SPC maintenance (Zhang et al., 1997; Crittenden et al., 2002). Despite 89% identity between FBF-1 and FBF-2 protein sequences, several reports suggest that FBF-1 and FBF-2 localize to distinct cytoplasmic RNA granules and have unique effects on the germline SPC pool (Lamont and others 2004; Voronina and others 2012). Specifically, FBF-1 and FBF-2 each support distinct numbers of SPCs (Lamont and others 2004). Furthermore, FBF-1 inhibits accumulation of target mRNAs in SPCs, while FBF-2 primarily represses translation of the target mRNAs (Voronina and others 2012). Some unique aspects of FBF-1 and FBF-2 function might be explained by their association with distinct protein cofactors, as we previously found that a small protein DLC-1 is a cofactor specific to FBF-2 that promotes FBF-2 localization and function (Wang and others 2016). Despite the fact that several repressive mechanisms have been documented for PUF family proteins (Quenault and others 2011), it is relatively understudied how the differences between PUF homologs are specified. Here we sought to take advantage of the distinct SPC numbers maintained by individual FBF proteins to understand how they regulate the dynamics of SPCs proliferation and differentiation and probe the functional differences between FBFs.

Elaborating on the general contribution of PUF proteins to stem cell maintenance, we describe here that FBF-1 and FBF-2 have opposing effects on the rate of germline SPCs proliferation and the rate of meiotic entry. We discovered that FBFs regulate core cell cycle machinery transcripts along with transcripts required for differentiation to coordinately change the steady-state amounts of both transcript classes. We show that FBF-1 decreases steady-state levels of target mRNAs and requires CCR4-NOT deadenylation machinery. By contrast, FBF-2 functions independently of CCR4-NOT and promotes accumulation of target mRNAs. These distinct functions of FBFs are determined by the protein regions outside of the conserved PUF homology domain. The dual regulation of SPC self-renewal and differentiation by FBFs effectively allows the stem cells to match cell division rate with the demand for meiotic cell output.

## RESULTS

### FBF-1 and FBF-2 differentially modulate proliferation and meiotic entry of *C. elegans* germline SPCs

During tissue maintenance, stem cells adjust their proliferative activity and differentiation rate to meet the physiological tissue demands through diverse regulatory mechanisms, including RNA-binding protein mediated post-transcriptional regulation. We hypothesized that two paralogous RNA-binding proteins FBF-1 and FBF-2 differentially regulate germline stem cell proliferation and differentiation in *C. elegans*, resulting in distinct effects on the size of stem and progenitor cell (SPC) zone. We first determined how the SPC zone size was affected by loss-of-function mutations of each *fbf*. SPCs were marked by staining for a nucleoplasmic marker REC-8 (**Figure 1A and C**) (Hansen and others 2004), and the SPC zone size was measured by counting the number of cell rows positive for REC-8 staining in each germline. Consistent with a previous report (Lamont and others 2004), we observed that the SPC zone of *fbf-1(ok91, loss-of-function mutation, lf)* (∼15 germ cell diameters, gcd; **Figure 1Ci**) is smaller than that of the wild type (∼20 gcd, **Figure 1Cii**), whereas the SPC zone of *fbf-2(q738, loss-of-function mutation, lf)* (∼25 gcd, **Figure 1Ciii**) is larger than that of the wild type (**Figures 1B and C**). The differences in SPC zone size between *fbf* single mutants and the wild type are consistently observed in animals from the late L4 to the second day of adulthood (**Figure 1--figure supplement 1A**).

To test whether the differences in germline SPC zone sizes between *fbf* mutants and the wild type result from changes in cell proliferation, we compared cell cycle parameters in each genetic background. We started with measuring the M-phase index (the percentage of SPC zone cells in M phase) following immunostaining for the SPC marker REC-8 and the M-phase marker phospho-histone H3 (pH3, **Figure 1C**). We found that the mitotic index of *fbf-1(lf)* was significantly higher than that of the wild type (by 54%, **Figure 1D**). By contrast, the mitotic index of *fbf-2(lf)* was significantly lower than that of the wild type (by 42%; **Figure 1D**). These results suggested that loss of FBF-1 might result in greater SPC proliferation, while loss of FBF-2 might reduce SPC proliferation. Since *C. elegans* stem cells have an abbreviated G1 and an extended G2 phases (Fox and others 2011), we tested whether the G2-phase duration is affected differentially by loss of function mutation of each *fbf.* Using phospho-histone H3 immunostaining and 5-ethynyl-2’-deoxyuridine (EdU) pulse we estimated a median G2 length by determining when 50% of pH3 positive cells become EdU-positive (**Figure 1—figure supplement 1B**). We found that the median G2 length of *fbf-2(lf)* is significantly greater than that of the wild type, suggesting that loss of FBF-2 results in slower progression through the G2-phase of the cell cycle (by 25%; **Figure 1E**). By contrast, the median G2 length of *fbf-1(lf)* is not significantly different from that of the wild type (**Figure 1E**). We conclude that FBF-2 promotes SPC proliferation by facilitating the G2-phase progression.

Despite an increase in mitotic index of *fbf-1(lf)*, its SPC zone is smaller than that of the wild type, suggesting a possibility that *fbf-1(lf)* might result in faster meiotic entry. Conversely, compared to the wild type, *fbf-2(lf)* maintains a relatively larger SPC population but with less proliferation, suggesting that the rate of meiotic entry in *fbf-2(lf)* might be slower than in the wild type. To test these possibilities, we determined the rate of meiotic entry in each genetic background. Animals were continuously EdU labeled and stained for EdU and REC-8 at three time points. The number of germ cells negative for REC-8 but positive for EdU were scored at each time point and the rate of meiotic entry was estimated from the slope of plotted regression line as in **Figure 1—figure supplement 1C**. We found that *fbf-1(lf)* results in a significantly increased rate of meiotic entry compared to the wild type (by 31%; **Figure 1F**), whereas *fbf-2(lf)* results in a significantly reduced rate of meiotic entry (by 18%; **Figure 1F**). We conclude that FBF-2 stimulates meiotic entry while FBF-1 inhibits meiotic entry.

In summary, mutations in *fbf-1* and *fbf-2* differentially influence both SPC proliferative activity and meiotic entry rate, suggesting FBF proteins have distinct effects on SPC proliferation and differentiation. FBF-1 promotes a more quiescent stem cell state characterized by a slower rate of meiotic entry, while FBF-2 promotes a more activated stem cell state characterized by faster rates of both cell cycle and meiotic entry.

### FBF-1 and FBF-2 differentially regulate mRNA abundance of target genes controlling proliferation and differentiation

FBFs are two redundant translational repressors in *C. elegans* germline SPCs. Although FBF-1 and FBF-2 share the majority of target mRNAs and bind to the same motif in the 3’UTRs (Porter and others 2019; Prasad and others 2016), they have different effects on their targets: FBF-1 promotes target mRNA clearance in the stem cell region, whereas FBF-2 sequesters target mRNAs (Voronina and others 2012). We hypothesized that the FBF-mediated effects on germline SPC proliferation and differentiation might be explained by their differential regulation of target mRNAs associated with proliferation and differentiation in germline SPCs. To test this hypothesis, we compared the steady-state mRNA abundance of selected FBF targets among the wild type, *fbf-1(lf)* and *fbf-2(lf)* genetic backgrounds by qPCR (**Figure 1G**). RNA samples were extracted from animals of *glp-1 (gain-of-function, gf)* mutant background, which produce germlines with only mitotic cells when grown at restrictive temperature, thus allowing us to focus on the changes in mRNA abundance in the mitotic cell population. We determined steady-state levels of meiotic entry associated transcripts, *him-3, htp-1,* and *htp-2* (previously described FBF targets (Merritt and Seydoux 2010)) and cell cycle regulators, *cyb-1, cyb-2.1, cyb-2.2* and *cyb-3* (FBF-bound transcripts (Kershner and Kimble 2010; Porter and others 2019; Prasad and others 2016)), as well as a control not regulated by FBFs, tubulin (*tbb-2*). All transcript levels were normalized to a housekeeping gene actin (*act-1*). We found that the mRNA levels of all tested FBF targets, except for *cyb-1*, are increased in *fbf-1(lf)* relative to the wild type and all are decreased in *fbf-2(lf)* relative to the wild type (**Figure 1G**). Linear trend analysis showed that the decrease in mRNA abundance of FBF targets from *fbf-1(lf)* to wild type to *fbf-2(lf)* is statistically significant (*P*<0.01); and the mRNA abundance of *htp-2* and all cyclin B genes among all three genetic backgrounds are significantly different by ANOVA analysis (*P*<0.01). The most dramatic change (5-fold difference) in mRNA abundance between *fbf-2(lf)* and *fbf-1(lf)* genetic backgrounds was observed for *cyb-2.1* mRNA. By contrast, the mRNA abundance of *tbb-2* control is not significantly different among the three analyzed genetic backgrounds (**Figure 1G**).

These findings suggest that FBF-1 might destabilize the target mRNAs controlling germline SPC proliferation and differentiation while FBF-2 promotes accumulation of the same target mRNAs. The distinct effects of the FBF homologs on target mRNAs may explain FBFs’ regulation of germline SPC proliferation and differentiation. For example, slower rates of cell cycle and meiotic entry in *fbf-2(lf)* genetic background might result from FBF-1-mediated destabilization of target mRNAs required for cell proliferation and differentiation. Next, we tested whether disrupting FBF-mediated regulation of a target transcript controlling cell cycle in *fbf-2(lf)* would influence the size of germline SPC zone.

### Repression of cyclin B by FBF limits accumulation of germline SPCs

Cyclin B/Cdk1 kinase, also known as M-phase promoting factor, triggers G2/M transition in most eukaryotes (Lindqvist and others 2009). Four cyclin B family genes provide overlapping as well as specific mitotic functions in *C. elegans* (van der Voet and others 2009). We hypothesized that the slower G2-phase and lower M-phase index of *fbf-2(lf)* SPCs results from FBF-1-mediated translational repression and reduced steady-state levels of four cyclin B family transcripts. We addressed this hypothesis in two ways. First, we tested whether mutation of <u>F</u>BF <u>b</u>inding <u>e</u>lements (FBEs) in the 3’UTR of *cyb-2.1* mRNA would result in translational derepression of *cyb-2.1*. Second, we assessed whether derepression of *cyb-2.1* in *fbf-2(lf)* would lead to accumulation of more SPCs due to greater proliferation but unchanged meiotic entry rate.

FBFs repress their target mRNAs by binding to the <u>F</u>BF-<u>b</u>inding <u>e</u>lements (FBEs; UGUxxxAU) in the 3’UTRs (Bernstein and others 2005; Crittenden and others 2002; Merritt and Seydoux 2010). Four mRNAs encoding Cyclin B family members co-purify with FBF proteins and contain predicted FBEs in their 3’UTRs (Porter and others 2019; Prasad and others 2016). Since *cyb-2.1* contains more canonical FBE sites than the other cyclin B transcripts and the mRNA abundance of *cyb-2.1* varies most dramatically between *fbf* mutants and the wild type (**Figure 1G**), we chose to analyze the translational regulation of *cyb-2.1*. If FBFs repress translation of *cyb-2.1* by binding to FBEs, mutation of FBEs would cause derepression of CYB-2.1 protein. To test this prediction, we established a transgenic animal *3xflag::cyb-2.1(fbm)*, expressing 3xFLAG::CYB-2.1 under the control of 3’UTR with mutated FBEs (ACAxxxAU); as a control, a transgenic animal expressing 3x*flag::cyb-2.1(wt)* with wild type FBEs was also established (**Figure 2A**). By immunoblotting, we found that the expression of 3xFLAG::CYB-2.1 protein was increased in *3xflag::cyb-2.1(fbm)* animals compared to *3xflag::cyb-2.1(wt)*, suggesting that mutation of FBEs caused translational derepression (**Figure 2B**). The protein levels of 3xFLAG::CYB-2.1wt might be too low to be detectable by western blot.

**Figure 2.**
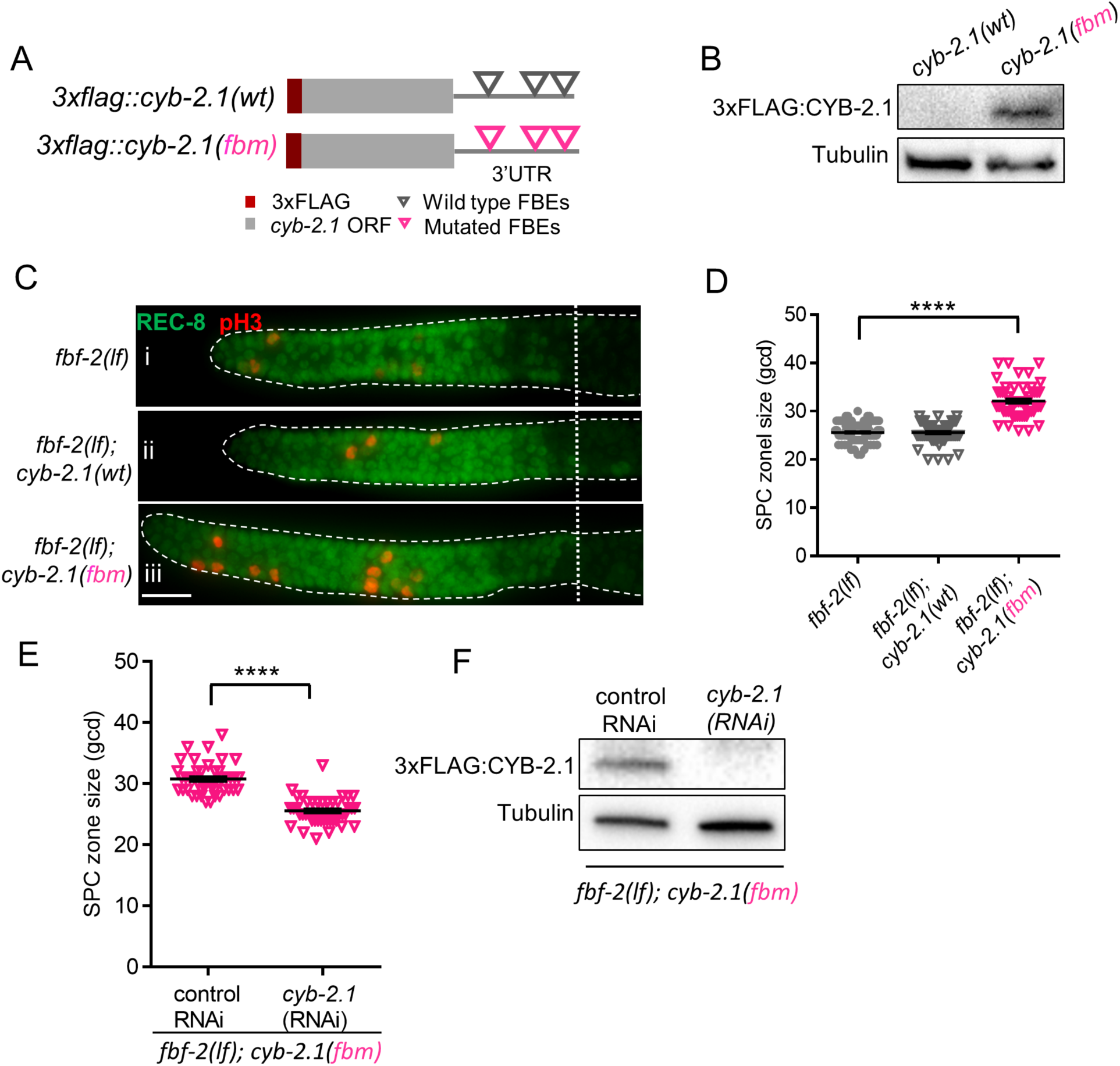
FBF-mediated repression of cyclin B limits accumulation of germline progenitor cells. (A) Schematic representation of transgenes encoding 3xFLAG-tagged CYB-2.1(wt) with wild type FBF binding elements (FBEs, UGUxxxAU) in 3’UTR and 3xFLAG-tagged CYB-2.1(fbm) with <u>F</u>BF <u>b</u>inding elements <u>m</u>utated (ACAxxxAU). (B) Immunoblot analysis of 3xFLAG::CYB-2.1 protein levels in *3xflag::cyb-2.1(wt)* and *3xflag::cyb-2.1(fbm)* worms using α-tubulin as a loading control. (C) Distal germlines dissected from the *fbf-2(lf), fbf-2(lf); cyb-2.1(fbm)* and *fbf-2(lf); cyb-2.1(wt)* animals and stained with anti-REC-8 (green) and anti-pH3 (red). Germlines are outlined with dashed lines and the vertical dotted line marks the beginning of transition zone. Scale bar: 10 μm. (D) Quantification of SPC zone size in the *fbf-2(lf), fbf-2(lf); cyb-2.1(fbm)* and *fbf-2(lf); cyb-2.1(wt)* genetic backgrounds. Plotted values are individual data points and arithmetical means ± S.E.M. Differences in SPC zone size were evaluated by one-way ANOVA with Dunnett’s post-test; asterisks mark statistically-significant difference (P<0.0001). Data was collected from 3 independent experiments and 57∼110 independent germlines were scored for each genotype. (E) Quantification of SPC zone size in the *fbf-2(lf); cyb-2.1(fbm)* after *cyb-2.1(RNAi)* compared to the empty vector RNAi control. Plotted values are individual data points and arithmetical means ± S.E.M. Differences in SPC zone size were evaluated by T-test; asterisks mark statistically-significant difference (*P*<0.0001). Data was collected from 2 independent experiments and 44 independent germlines were scored for each condition. (F) Immunoblot analysis of 3xFLAG::CYB-2.1 protein levels in *3xflag::cyb-2.1fbm* after *cyb-2.1(RNAi)* compared to the empty vector RNAi control. Tubulin was used as a loading control.

A larger SPC zone size in *fbf-2(lf)* is associated with slower SPC proliferation in conjunction with a slower SPC meiotic entry rate. We hypothesized that the slower SPC proliferation is caused by FBF-1-mediated destabilization and repression of cyclin B-family mRNAs. If any cyclin B-family gene can promote SPC proliferation, disrupting translational repression of a single cyclin B-family transcript in *fbf-2(lf)* would facilitate SPC proliferation, resulting in accumulation of SPCs and an increase of SPC zone size when SPC meiotic entry rate is unchanged. To test this hypothesis, we measured the SPC zone size after crossing the *3xflag::cyb-2.1fbm* and *3xflag::cyb-2.1wt* transgenes into *fbf-2(lf)* genetic background. We found that the SPC zone of *fbf-2(lf)*; *3xflag::cyb-2.1fbm* (∼32 gcd, **Figure 2Ciii**) is significantly larger than that of the *fbf-2(lf)* (∼26 gcd, **Figure 2Ci, D**, *P*< 0.0001). By contrast, there is no significant difference in the SPC zone size between the *fbf-2(lf)*; *3xflag::cyb-2.1wt* and *fbf-2(lf)* (**Figure 2Cii and D**). To test whether the expansion of SPC zone in *fbf-2(lf)*; *3xflag::cyb-2.1fbm* results from overexpression of *cyb-2.1*, we measured the SPC zone size following knockdown of *cyb-2.1* by RNAi. We found that the SPC zone of *fbf-2(lf); 3xflag::cyb-2.1fbm* after *cyb-2.1(RNAi)* became significantly smaller (∼ 26 gcd) compared to the control RNAi (∼31 gcd; **Figure 2E**). Depletion of CYB-2.1 was confirmed by immunoblot for FLAG::CYB-2.1 after RNAi of *cyb-2.1* compared to the control (**Figure 2F**).

We conclude that the levels of B-type cyclins limit SPC proliferation rate in *fbf-2(lf)* and disruption of FBF-1-mediated repression of a single cyclin B gene is sufficient to affect the size of germline SPC zone. We next focus on investigating the mechanism of FBF-1-mediated mRNA regulation.

### FBF-1 function requires CCR4-NOT deadenylase complex

One mechanism of PUF-dependent destabilization of target mRNAs is through recruitment of CCR4-NOT deadenylase that shortens poly(A) tails of the targets (Quenault and others 2011). CCR4-NOT deadenylase is a complex that includes three core subunits: two catalytic subunits CCR-4/CNOT6/6L and CCF-1/CNOT-7/8 and one scaffold subunit LET-711/CNOT1, which are highly conserved in *C. elegans* and humans (**Figure 3A**; (Nousch and others 2013). Although multiple PUF family proteins, including FBF homologs in *C. elegans*, interact with a catalytic subunit of CCR4-NOT *in vitro,* the contribution of CCR4-NOT to PUF-mediated repression *in vivo* is still controversial (Suh and others 2009; Weidmann and others 2014). We hypothesized that the enlarged germline SPC zone in *fbf-2(lf)* mutant results from FBF-1-mediated destabilization and translational repression of target mRNAs achieved through the activity of CCR4-NOT deadenylase. If so, knockdown of CCR4-NOT in *fbf-2(lf)* genetic background would lead to derepression of target mRNAs in SPCs and a decrease of SPC zone size.

**Figure 3.**
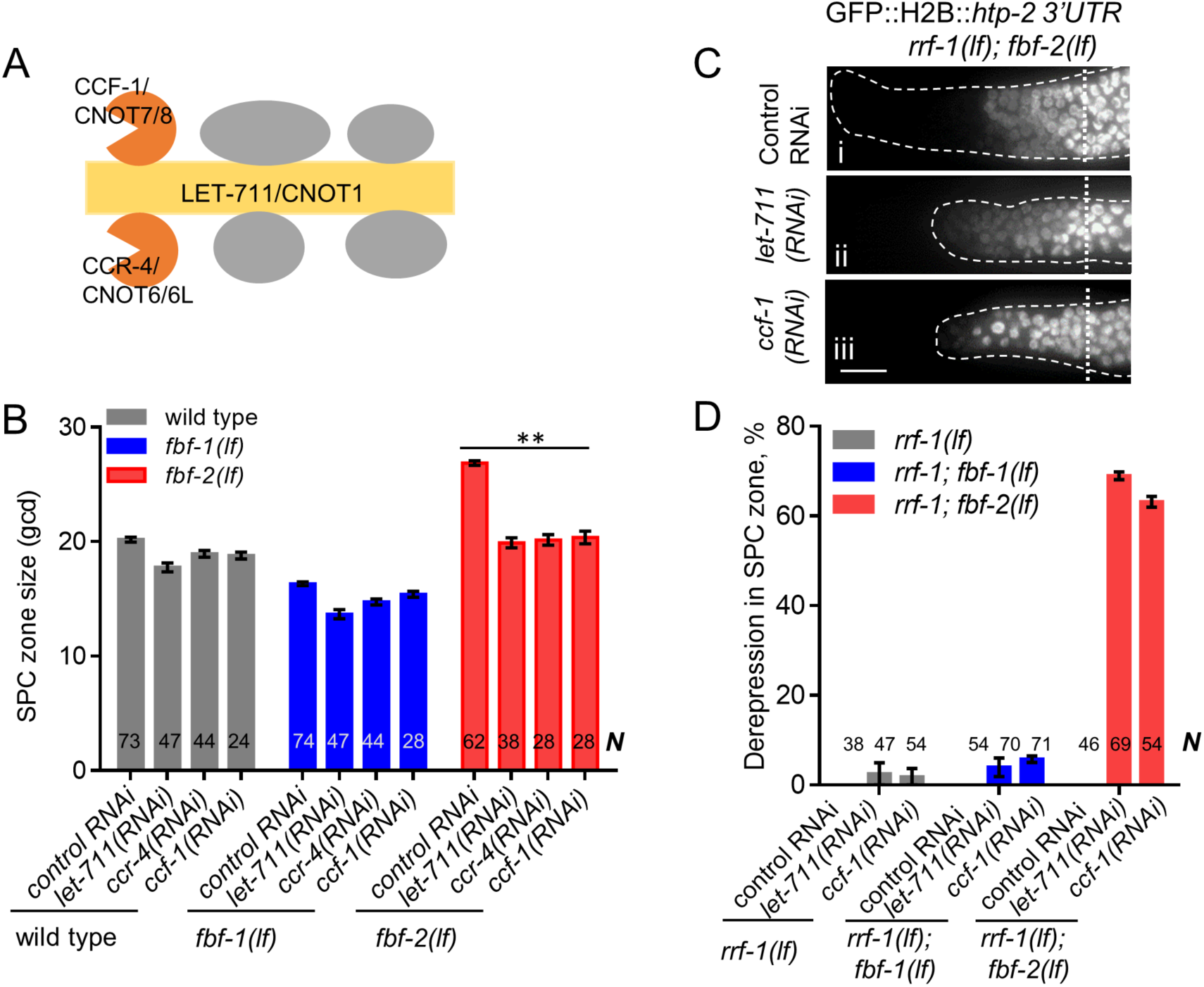
CCR4-NOT deadenylase complex promotes FBF-1 function in germline SPCs. (A) Schematic of CCR4-NOT deadenylase complex in humans and *C. elegans*. (B) Quantification of SPC zone size after knocking down CCR4-NOT subunits in the wild type, *fbf-1(lf)* and *fbf-2(lf)* genetic backgrounds. Genetic backgrounds and RNAi treatments are indicated on the X-axis and the average size of SPC zone ± S.E.M is plotted on the Y-axis. Differences in SPC zone size between CCR4-NOT RNAi and the empty vector RNAi control were evaluated by one-way ANOVA. Asterisks mark the group with significant changes in SPC zone sizes after CCR4-NOT knockdown, *P*<0.01. Data was collected from 3 independent experiments. N, the number of hermaphrodite germlines scored in each RNAi treatment. (C) Distal germlines of *rrf-1(lf); fbf-2(lf)* expressing a GFP::Histone H2B fusion under the control of the *htp-2* 3’UTR after the indicated RNAi treatments. Germlines are outlined with dashed lines and vertical dotted lines indicate the beginning of the transition zone. All images were taken with a standard exposure. Scale bar: 10 μm. (D) Percentage of germlines showing expression of GFP::H2B fusion extended to the distal end in the indicated genetic backgrounds and knockdown conditions. Plotted values are arithmetical means ± S.E.M. Data was collected from 3 independent experiments. *N*, the number of germlines scored. Efficiencies of RNAi treatments were confirmed by sterility (Figure 3—figure supplement 1B) or embryonic lethality (Supplemental Table 3).

First, we measured SPC zone size after RNAi-mediated knockdown of core CCR4-NOT subunits, and we found that CCR4-NOT RNAi dramatically shortened the SPC zone in *fbf-2(lf)* compared to the control RNAi (*P*<0.01; **Figure 3B**). By contrast, the sizes of SPC zones in the wild type and *fbf-1(lf)* animals were not significantly affected by CCR4-NOT knockdown (**Figure 3B**). These findings suggest that CCR4-NOT is required for FBF-1-mediated regulation of germline SPC zone size, but does not significantly contribute to FBF-2 function.

Next, we tested whether CCR4-NOT knockdown disrupts FBF-1-mediated translational repression in SPCs. One relevant FBF target mRNA is *htp-2*, a HORMA domain meiotic protein (Merritt and Seydoux 2010). Translational regulation of a transgenic reporter encoding GFP::Histone H2B fusion under the control of *htp-2* 3’UTR recapitulates FBF-mediated repression in germline SPCs (Merritt and Seydoux 2010). We performed CCR4-NOT RNAi in the *rrf-1(lf)* background to preferentially direct the RNAi effects to the germline and avoid any defects in the somatic cells (Kumsta and Hansen 2012; Sijen and others 2001) and observed derepression of the reporter in SPCs of 63-69% germlines of *rrf-1(lf); fbf-2(lf)* genetic background (**Figure 3C and D**). By contrast, derepression of the reporter was observed only in 3-5% of *rrf-1(lf)* and *rrf-1(lf); fbf-1(lf)* genetic backgrounds (**Figure 3D; Figure 3—figure supplement 1A**). These data suggest that the CCR4-NOT deadenylase complex is necessary for FBF-1-mediated translational repression of targets in germline SPCs, but is dispensable for FBF-2 regulatory function. In addition, we observed significantly increased sterility upon CCR4-NOT knockdown in *rrf-1(lf); fbf-2(lf)* compared to the *rrf-1(lf)* and *rrf-1(lf); fbf-1(lf)* (**Figure 3—figure supplement 1B**).

CCR4-NOT knockdown might disrupt FBF-1 regulatory function or FBF-1 protein expression and localization. To distinguish between these possibilities, we determined the abundance of endogenous FBF-1 after *ccf-1(RNAi)* by immunoblotting using tubulin as a loading control. We found that FBF-1 protein abundance is not decreased after CCF-1 knockdown compared to the control (**Figure 3—figure supplement 1C and D**). Immunostaining for the endogenous FBF-1 showed that in control germlines FBF-1 localized in foci adjacent to perinuclear P granules (**Figure 3—figure supplement 1E**) as previously reported (Voronina and others 2012). Upon CCF-1 knockdown, FBF-1 foci were still observed next to P granules (**Figure 3—figure supplement 1F**). Therefore, we conclude that CCR4-NOT is not required for FBF-1 expression and localization, and CCR4-NOT knockdown specifically disrupts FBF-1 function.

In summary, we conclude that CCR4-NOT is required for FBF-1, but not FBF-2-mediated regulation of target mRNA and germline SPC zone size. We further predicted that FBF-1 localizes together with CCR4-NOT to the same RNA-protein complex in SPCs.

### FBF-1 colocalizes with CCR4-NOT in germline SPCs

Using co-immunostaining of endogenous FBF-1 or GFP::FBF-1 and 3xFLAG::CCF-1 followed by Pearson’s correlation coefficient analysis based on Costes’ automatic threshold (Costes and others 2004), we found that both endogenous FBF-1 and GFP::FBF-1 foci colocalize with 3xFLAG::CCF-1 foci in SPC cytoplasm (**Figure 4A and C; Figure 4—figure supplement 1A and B**). By contrast, GFP::FBF-2 and 3xFLAG::CCF-1 do not colocalize (**Figure 4B and C**). As an alternative metric of colocalization, we used proximity ligation assay (PLA) that can detect protein-protein interactions *in situ* at the distances <40 nm (Fredriksson and others 2002). PLA was performed in *3xflag::ccf-1; gfp::fbf-1*, *3xflag::ccf-1; gfp::fbf-2*, and *3xflag::ccf-1; gfp* animals using the same antibodies and conditions for all three protein pairs. We observed significantly more dense PLA signals in *3xflag::ccf-1; gfp::fbf-1* than in the control (**Figure 4D**; p<0.0001, **Table 1**). By contrast, PLA foci density in mitotic germ cells of *3xflag::ccf-1; gfp::fbf-2* was not different from the control (**Figure 4D; Table 1**), although the expression of GFP::FBFs or GFP alone in mitotic germ cells appeared similar (**Figure 4—figure supplement 1C**). Together, these data suggest that FBF-1, but not FBF-2, colocalizes with CCR4-NOT in SPCs, in agreement with the dependence of FBF-1 function on CCR4-NOT.

**Figure 4.**
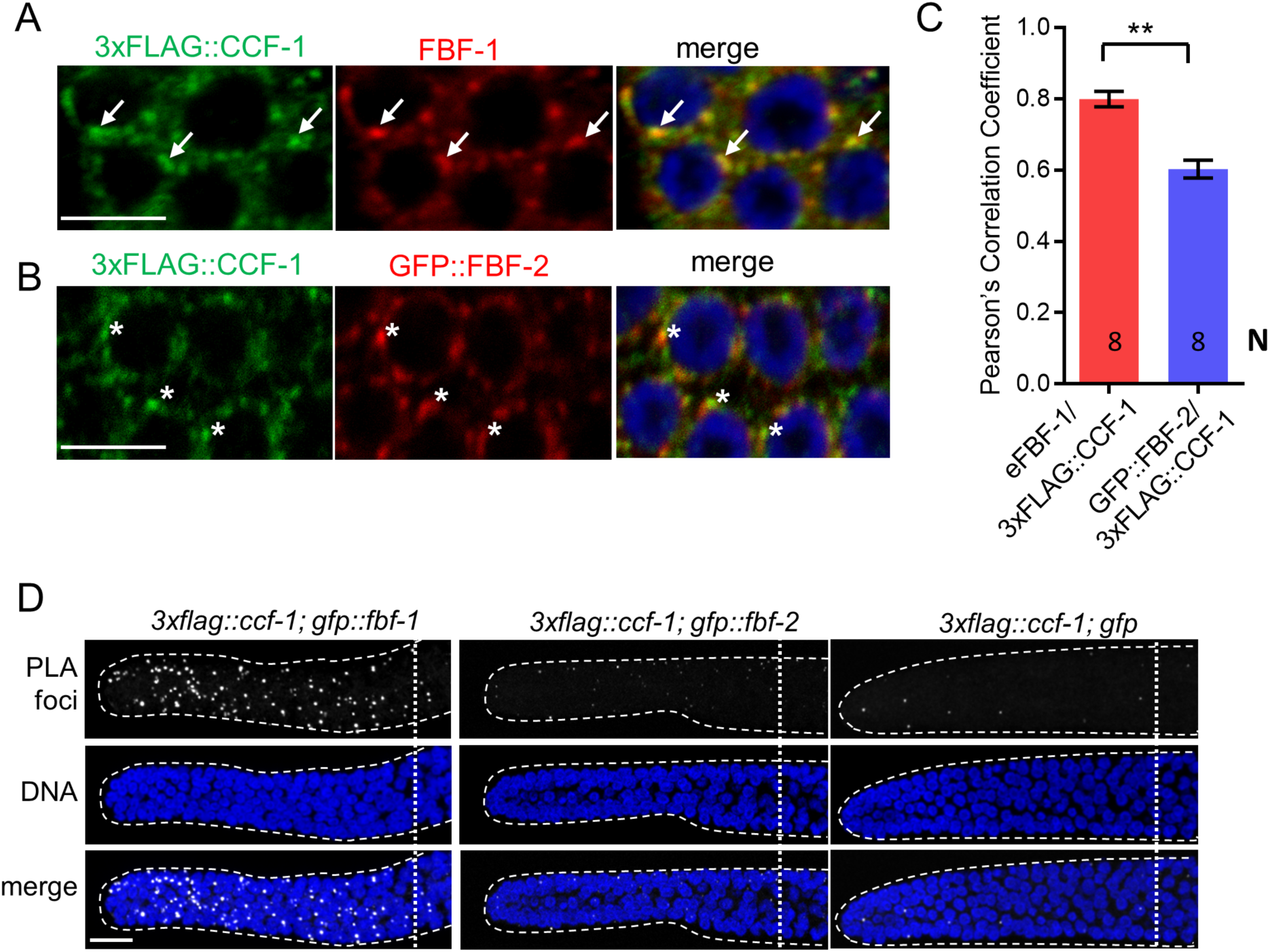
FBF-1 colocalizes with CCR4-NOT complex in germline SPCs. (A-B) Confocal images of SPCs co-immunostained for endogenous FBF-1 (A) or GFP-tagged FBF-2 (B, red) and 3xFLAG-tagged CCF-1 (green). DNA staining is in blue (DAPI). Arrows indicate complete overlap of FBF-1 and CCF-1 granules. Asterisks denote FBF-2 granules localizing close but not overlapping with CCF-1 granules. Scale bars in A and B: 5 μm. (C) Pearson’s correlation analysis quantifying the colocalization between FBF and CCF-1 granules in co-stained germline images. Plotted values are arithmetical means ± S.E.M. *N*, the number of analyzed germline images (single confocal sections through the middle of germline SPC nuclei including 5-8 germ cells). Statistical analysis was performed by Student’s t-test, asterisks mark statistically significant difference, *P*<0.01. (D) Confocal images of the distal germline SPC zones with PLA foci (grayscale) and DNA staining (blue). Germlines are outlined with dashed lines and vertical dotted lines indicate the beginning of the transition zone. Genotypes are indicated on top of each image group. Scale bar: 10 μm.

**Table 1.**
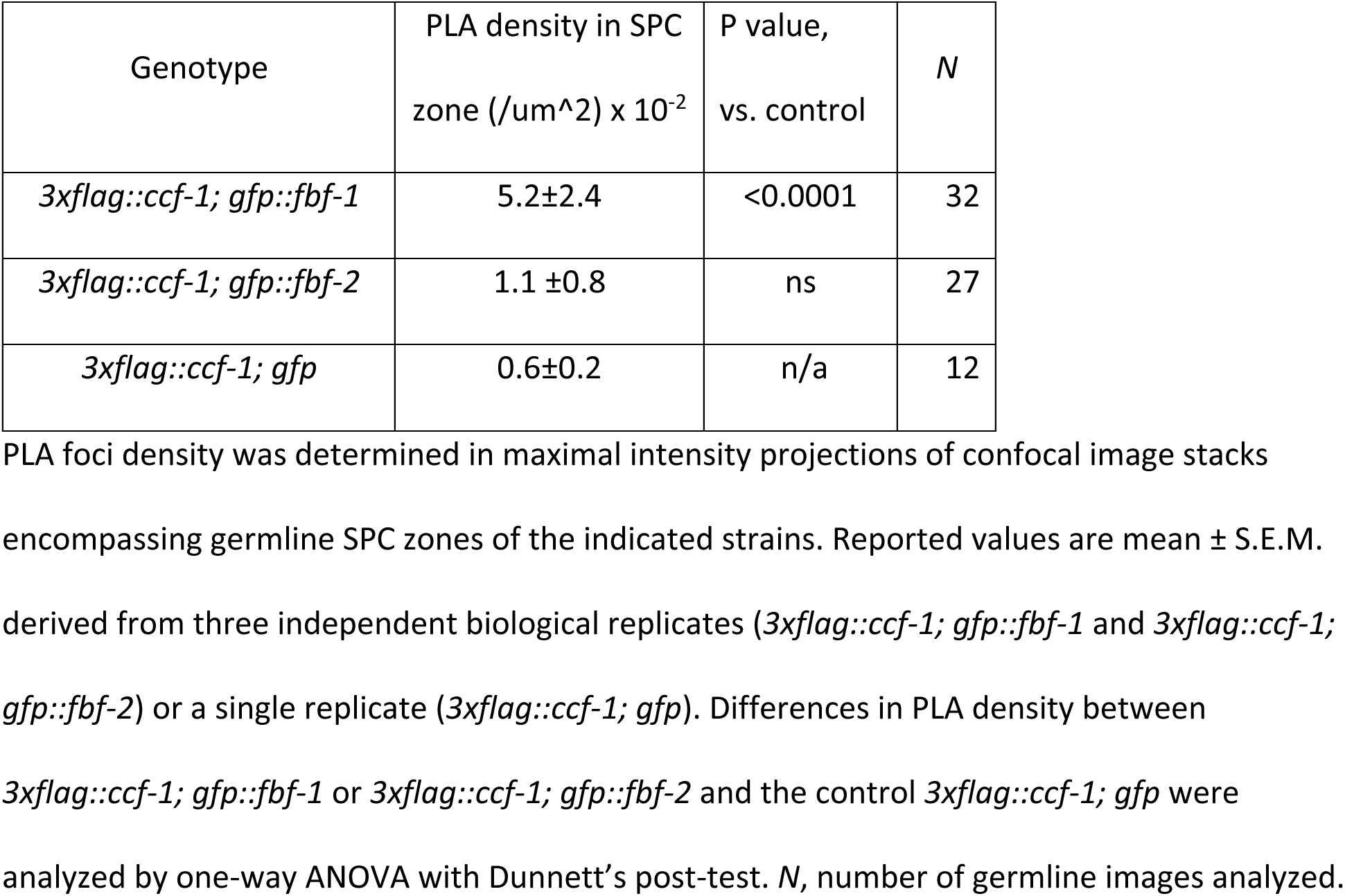
Proximity ligation assay density analysis

### FBF-1 promotes deadenylation of its target mRNA

Since a knockdown of CCR4-NOT deadenylase compromises FBF-1-mediated target repression, we hypothesized that FBF-1 promotes deadenylation of target mRNAs. We investigated whether the lower abundance of *cyb-2.1* mRNA in *fbf-2(lf)* correlated with a shorter poly(A) tail length. Poly(A) tail (PAT)-PCR for *cyb-2.1* and control *tbb-2* were performed to determine the poly(A) tail length using RNA samples extracted from *fbf-1(lf); glp-1(gf)* and *fbf-2(lf); glp-1(gf)*. PAT-PCR assays revealed that the poly(A) tail length of the predominant *cyb-2.1* mRNA species in *fbf-2(lf)* is shorter than that in *fbf-1(lf)* (**Figure 5A and C**). By contrast, the poly(A) tail lengths of *tbb-2* tubulin mRNA in *fbf-2(lf)* and *fbf-1(lf)* are similar (**Figure 5B and D**). We conclude that FBF-1 promotes deadenylation of its target mRNAs.

**Figure 5.**
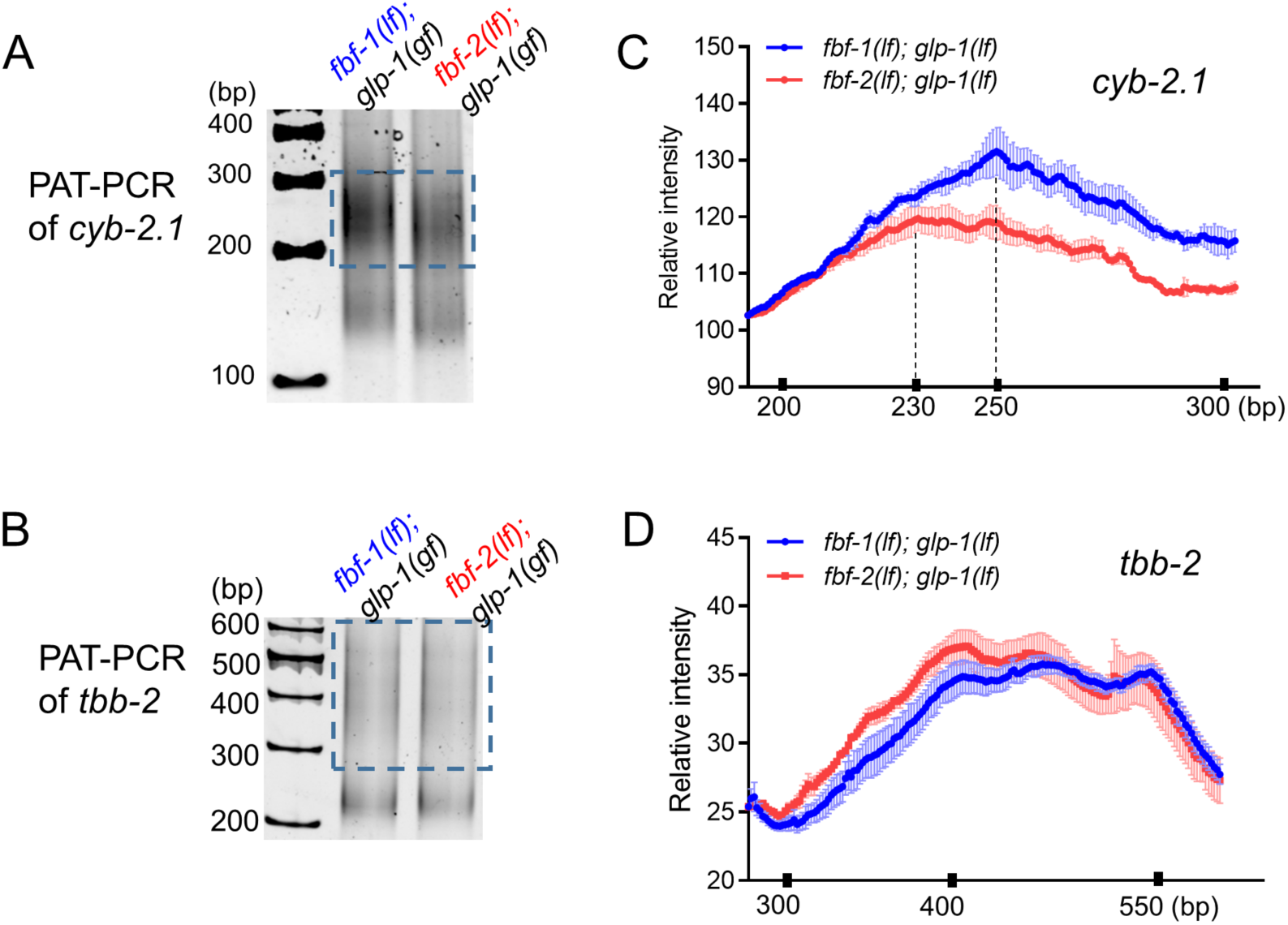
FBF-1 promotes deadenylation of *cyb-2.1* mRNA. (A, B) Representative PAT-PCR analysis of the poly(A) tail length of *cyb-2.1* mRNA (A) and control *tbb-2* mRNA (B) in *fbf-1(lf); glp-1(gf)* and *fbf-2(lf); glp-1(gf)* genetic backgrounds. The positions of size markers are indicated on the left. The areas boxed by dotted lines were quantified by densitometry in ImageJ. (C, D) Densitometric quantification of the *cyb-2.1* and *tbb-2* PAT-PCR amplification products (boxed in A, B). Y-axis, the relative intensity (arbitrary units) representing the average of PAT-PCR reactions from three independent biological replicates. X-axis, sizes of analyzed polyadenylated mRNA species. Values are arithmetical means ± S.E.M. Vertical dashed lines in (C) mark the sizes of the most abundant species of polyadenylated *cyb-2.1* mRNA in each *fbf* mutant background.

### Three variable regions outside of FBF-2 RNA binding domain are necessary to prevent cooperation with CCR4-NOT

Our findings suggest that FBF-1-mediated SPC maintenance depends on CCR4-NOT deadenylase complex, while FBF-2 can function independent of CCR4-NOT. Since FBF proteins are very similar in primary sequence except for the four <u>v</u>ariable <u>r</u>egions (VRs, **Figure 6A**), we next investigated whether the VRs were necessary for FBF-2-specific maintenance of germline SPCs and prevented FBF-2 dependence on CCR4-NOT. We previously found that mutations/deletions of the VRs outside of FBF-2 RNA-binding domain (VR1, 2 and 4, **Figure 6A**) produced GFP::FBF-2(vrm) protein with a disrupted localization and compromised function (Wang and others 2016). We hypothesized that these three VRs might contribute to FBF-2-specific effects on SPC zone size as well as prevent FBF-2 from cooperating with CCR4-NOT.

**Figure 6.**
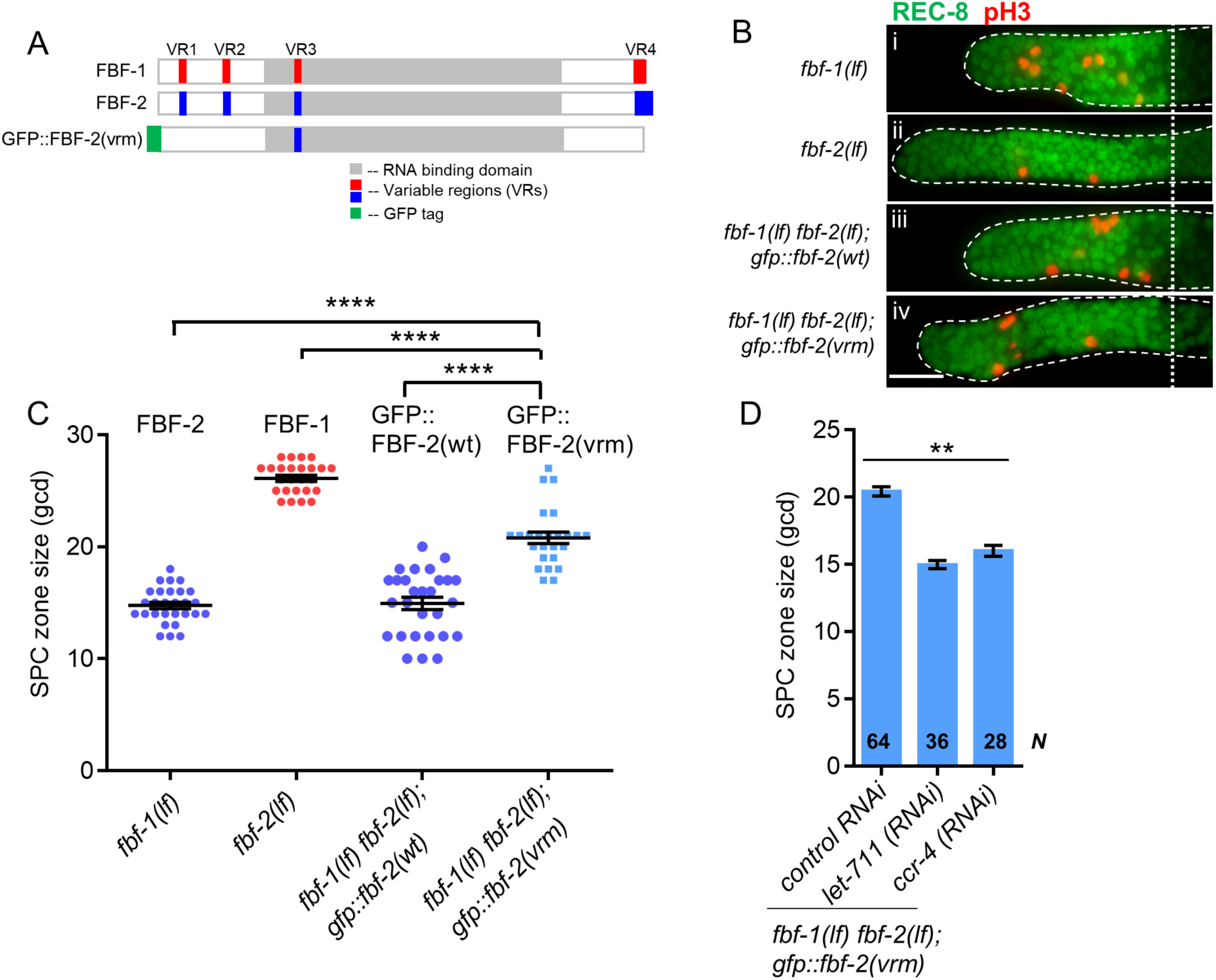
Three variable regions of FBF-2 prevent its cooperation with CCR4-NOT. (A) Schematics of FBF-1, FBF-2 and GFP::FBF-2(vrm) mutant transgene (Wang and others 2016). Red and blue colors indicate variable regions distinguishing FBF-1 and FBF-2 respectively, grey box indicates the RNA binding domain, and green box indicates GFP tag. (B) Distal germlines of the indicated genetic backgrounds stained with anti-REC-8 (green) and anti-pH3 (red). Germlines are outlined with the dashed lines, and the vertical dotted line marks the beginning of transition zone. Scale bar: 10 μm. (C) Germline SPC zone sizes in the indicated genetic backgrounds (indicated on the X-axis). FBF protein(s) present in each genetic background are indicated above each data set. Plotted values are individual data points and arithmetical means ± S.E.M. Differences in SPC zone size between *fbf-1(lf) fbf-2(lf); gfp::fbf-2(vrm)* and the other strains were evaluated by one-way ANOVA test with Dunnett’s post-test; asterisks mark statistically significant differences (P<0.0001). Data were collected from 3 independent experiments and 24-28 germlines were scored for each genotype. (D) Quantification of SPC zone size after knocking down CCR4-NOT subunits in the *fbf-1(lf) fbf-2(lf); gfp::fbf-2(vrm)* genetic background. RNAi treatments are indicated on the X-axis and average size of SPC zone ± S.E.M on the Y-axis. Differences in SPC zone sizes between CCR4-NOT knockdowns and control were evaluated by one-way ANOVA (asterisks, *P*<0.01). Data were collected from 3 independent experiments. *N*, the number of independent germlines scored.

We first tested whether the three VRs are required for FBF-2-specific SPC zone size. To test this hypothesis, SPC zone size was determined after crossing the GFP::FBF-2(vrm) transgene into *fbf* double mutant background. We found that the SPC zone size maintained by GFP::FBF-2(vrm) (**Figure 6Bv**) is significantly larger than that maintained by GFP::FBF-2(wt) (**Figure 6Biv**) and the endogenous FBF-2 (**Figure 6Bii**) and significantly shorter than that maintained by FBF-1 (P<0.01, **Figure 6C**), suggesting that the GFP::FBF-2(vrm) effect on SPC zone size is distinct from that of FBF-2. To test whether the GFP::FBF-2(vrm) can rescue either of *fbf* single mutants, we determined the SPC zone size after crossing GFP::FBF-2(vrm) into *fbf-1(lf)* and *fbf-2(lf)* genetic backgrounds. As controls, the size of SPC zones were also measured after crossing the wild type GFP::FBF-2(wt) and GFP::FBF-1(wt) transgenes into each *fbf* single mutant. As expected, the SPC zone size of *fbf-2(lf); gfp::fbf-2(wt)* is significantly smaller than *fbf-2(lf)* (P<0.01) while the SPC zone size of *fbf-2(lf); gfp::fbf-1(wt)* is similar to *fbf-2(lf)* (**Figure 6—figure supplement 1A**), suggesting that GFP::FBF-2(wt), but not GFP::FBF-1(wt), rescues *fbf-2(lf)*. Likewise, GFP::FBF-1(wt), but not GFP::FBF-2(wt), rescues *fbf-1(lf)* (P<0.01, **Figure 6—figure supplement 1B**).

Interestingly, we found that the SPC zone size of *fbf-2(lf); gfp::fbf-2(vrm)* is similar to that of *fbf-2(lf)* (**Figure 6—figure supplement 1A**), suggesting that GFP::FBF-2(vrm) does not rescue *fbf-2(lf)*. By contrast, the SPC zone of *fbf-1(lf); gfp::fbf-2(vrm)* is significantly larger than that of *fbf-1(lf)* (P<0.01, **Figure 6—figure supplement 1B**) and there is no significant difference in the SPC zone between *fbf-1(lf); gfp::fbf-2(vrm)* and the wild type, suggesting that the GFP::FBF-2(vrm) completely rescues *fbf-1(lf).* We conclude that the three VRs outside of FBF-2 RNA-binding domain (VR1, 2, and 4) are important for FBF-2-specific effect on germline SPC zone size and mutation or deletion of these VRs resulted in a mutant protein FBF-2(vrm) that functions similar to FBF-1.

Since FBF-1 function requires CCR4-NOT complex and FBF-2(vrm) appears similar to FBF-1, we hypothesized that CCR4-NOT is required for FBF-2(vrm)-mediated function. To test this hypothesis, we measured SPC zone size after knockdown of CCR4-NOT subunits in *fbf-1(lf) fbf-2(lf); gfp::fbf-2(vrm)* animals by RNAi. We found that SPC zone size of *fbf-1(lf) fbf-2(lf); gfp::fbf-2(vrm)* after RNAi of CCR4-NOT subunits becomes significantly shorter than the control (P<0.01, **Figure 6D**), suggesting that GFP::FBF-2(vrm) function requires CCR4-NOT. We conclude that the VRs outside of FBF-2 RNA-binding domain are required for FBF-2-specific effect on SPC zone size and to prevent FBF-2 from cooperation with CCR4-NOT.

### The variable region 4 (VR4) of FBF-2 is sufficient to prevent cooperation with CCR4-NOT

To test whether one of the three VRs outside of FBF-2 RNA-binding domain (VR1, 2, and 4) is sufficient to support FBF-2-specific effects on SPC zone size, we established a transgenic FBF-1 chimera with VR4 swapped from FBF-2 (GFP::FBF-1(vr4sw); **Figure 7A**) and crossed it into *fbf* double mutant. Since VR3 residing in FBF-2 RNA-binding domain was not sufficient for FBF-2-specific function, *fbf-1(lf) fbf-2(lf); gfp::fbf-1(vr3sw)* (with VR3 swapped from FBF-2; **Figure 7A**) chimeric transgene was made for comparison. SPC zone size assessment showed that the SPC zone maintained by GFP::FBF-1(vr4sw) (**Figure 7Biii**) is significantly smaller than that maintained by GFP::FBF-1(wt) (**Figure 7Bv**) and endogenous FBF-1 (P<0.0001; **Figure 7Bii and C**). By contrast, the SPC zone maintained by GFP::FBF-1(vr3sw) (**Figure 7Biv**) is similar to that maintained by the GFP::FBF-1(wt) (**Figure 7Biv and C**). This finding suggested that GFP::FBF-1(vr4sw) might function similarly to FBF-2. To test whether GFP::FBF-1(vr4sw) rescues FBF-1- or FBF-2-specific function, we measured the sizes of SPC zones after crossing GFP::FBF-1(vr4sw) into *fbf-1(lf)* and *fbf-2(lf)* genetic backgrounds. For comparison, GFP::FBF-1(vr3sw) was also crossed into each *fbf* single mutant. We found that the SPC zone size of *fbf-1(lf); gfp::fbf-1(vr4sw)* is similar to that of *fbf-1(lf)* (**Figure 7—figure supplement 1A**), suggesting that GFP::FBF-1(vr4sw) does not rescue *fbf-1(lf)*. Interestingly, SPC zone size of *fbf-2(lf); gfp::fbf-1(vr4sw)* is significantly smaller than that of *fbf-2(lf)* (*P*<0.01, **Figure 7—figure supplement 1B**), suggesting that GFP::FBF-1(vr4sw) rescues *fbf-2(lf)*. By contrast, GFP::FBF-1(vr3sw) rescues *fbf-1(lf),* but not *fbf-2(lf)* (**Figure 7—figure supplement 1A and B**). We conclude that the presence of VR4 from FBF-2 in a chimeric GFP::FBF-1(vr4sw) protein is sufficient to impart FBF-2-specific effect on SPC zone size.

**Figure 7.**
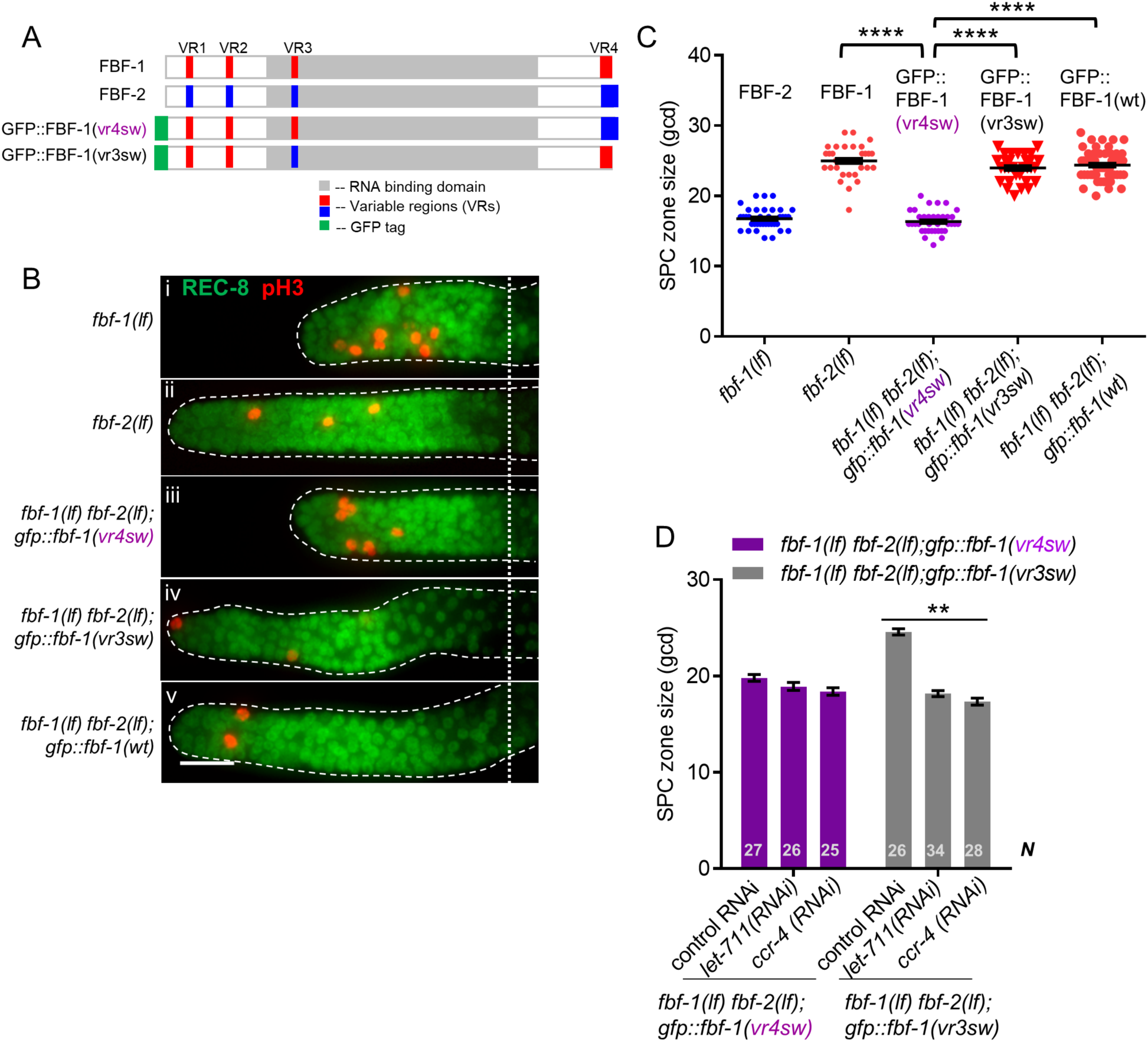
Variable region 4 (VR4) from FBF-2 is sufficient to prevent FBF-1 chimera from cooperation with CCR4-NOT. (A) Schematics of FBF-1, FBF-2, transgenic GFP::FBF-1(vr4sw) chimera (with VR4 swapped from FBF-2) and transgenic GFP::FBF-1(vr3sw) chimera (with VR3 swapped from FBF-2). Red and blue colors indicate variable regions distinguishing FBF-1 and FBF-2 respectively, grey box indicates RNA binding domain, and green box indicates GFP tag. (B) Distal germlines dissected from the indicated genetic backgrounds stained with anti-REC-8 (green) and anti-pH3 (red). Germlines are outlined with the dashed lines and the vertical dotted line marks the beginning of the transition zone. Scale bar: 10 μm. (C) Germline SPC zone sizes in the indicated genetic backgrounds (indicated on the X-axis). FBF protein present in each genetic background is indicated above each data set. Plotted values are individual data points and arithmetical means ± S.E.M. Differences in SPC zone sizes between *fbf-1(lf) fbf-2(lf); gfp::fbf-1(vr4sw)* and the other strains were evaluated by one-way ANOVA test with Dunnett’s post-test; asterisks mark statistically significant differences (*P*<0.0001). Data were collected from 2 independent experiments and 31-60 germlines were scored for each genotype. (D) Quantification of SPC zone size after knocking down CCR4-NOT subunits in the *fbf-1(lf) fbf-2(lf); gfp::fbf-1(vr4sw)* and *fbf-1(lf) fbf-2(lf); gfp::fbf-1(vr3sw)* genetic backgrounds (indicated on the X-axis). Plotted values are arithmetical means ± S.E.M. Differences in SPC zone sizes between CCR4-NOT RNAi and control RNAi were evaluated by one-way ANOVA. Asterisks mark the group with significant changes in SPC zone sizes after CCR4-NOT knockdown (*P*<0.01). Data was collected from 2 independent experiments. *N*, the number of hermaphrodite germlines scored.

To test whether VR4 is sufficient to inhibit cooperation of GFP::FBF-1(vr4sw) with CCR4-NOT, we measured the size of SPC zone after knockdown of CCR4-NOT subunits in *fbf-1(lf) fbf-2(lf); gfp::fbf-1(vr4sw)* animals by RNAi. As a control, CCR4-NOT knockdown was also performed on *fbf-1(lf) fbf-2(lf); gfp::fbf-1(vr3sw).* We found that the SPC zone of *fbf-1(lf) fbf-2(lf); gfp::fbf-1(vr4sw)* after RNAi of CCR4-NOT subunits is similar to the control (**Figure 7D**), suggesting that GFP::FBF-1(vr4sw) function in SPCs does not rely on CCR4-NOT. By contrast, the SPC zone of *fbf-1(lf) fbf-2(lf); gfp::fbf-1(vr3sw)* is significantly shortened after RNAi of CCR4-NOT subunits compared to the control (*P*<0.01, **Figure 7D**), indicating that GFP::FBF-1(vr3sw) maintains dependence on CCR4-NOT. We conclude that FBF-2 VR4 in a chimeric GFP::FBF-1(vr4sw) protein is sufficient to support FBF-2-specific effect on germline SPC zone size and to prevent the chimera’s cooperation with CCR4-NOT.

## DISCUSSION

This manuscript focuses on the roles of PUF family FBF proteins in the control of proliferation and differentiation of *C. elegans* germline stem and progenitor cells. Our results support three main conclusions. First, FBF proteins affect SPC proliferation and differentiation through translational control of FBF target mRNAs required for both processes. Second, FBF-mediated repression of cyclin B affects SPC proliferation. Third, distinct effects of FBF homologs on SPC development and their target mRNAs are mediated by differential cooperation of FBFs with deadenylation machinery. In turn, activation of deadenylation machinery by FBFs depends on the protein sequences outside of the conserved PUF RNA-binding domain. Collectively, our results support a model where the output of stem cell population is controlled by two paralogous proteins that have complementary effects on SPC proliferation and differentiation achieved through distinct regulatory mechanisms (**Figure 8**).

**Figure 8.**
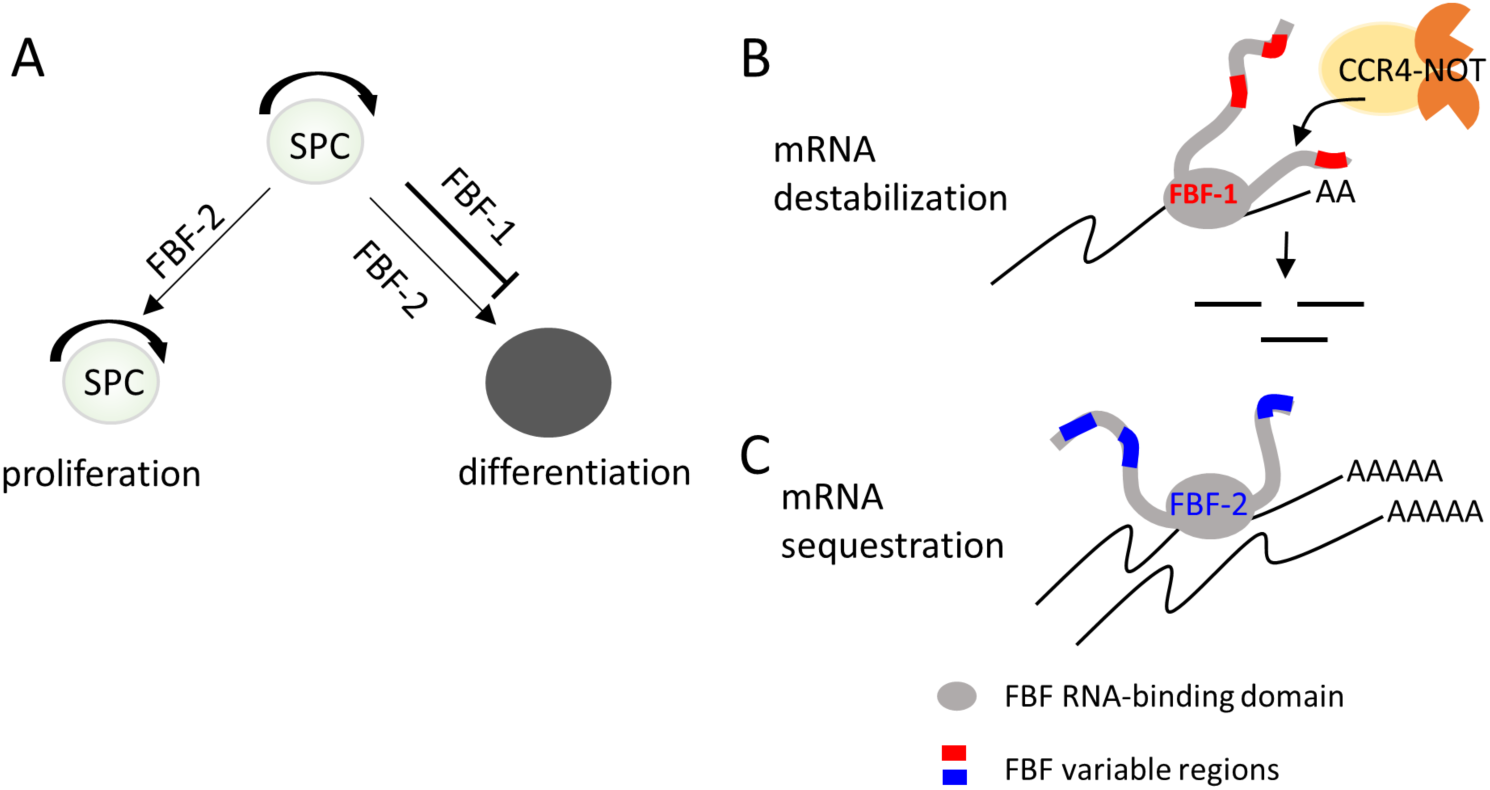
Distinct effects of FBF-1 and FBF-2 on germline SPC dynamics are mediated by their effects on target mRNAs in *C. elegans*. (A) Complementary activities of FBFs in maintaining germline SPC homeostasis: FBF-1 promotes SPC self-renewal by inhibiting differentiation, while FBF-2 facilitates both proliferation and differentiation of SPCs. (B, C) FBFs differentially control target mRNAs that regulate both stem cell proliferation and differentiation, and FBFs differential cooperation with CCR4-NOT is determined by their variable regions, VRs, outside of the RNA binding domain of FBFs. (B) FBF-1 cooperates with CCR4-NOT deadenylase and destabilizes target mRNAs. FBF-1-dependent RNA regulation is required to restrict the rate of germline stem cell differentiation. (C) FBF-2 does not rely on CCR4-NOT and promotes accumulation of target mRNAs. FBF-2-dependent accumulation of mRNAs is required to sustain both wild type rates of germline stem cell proliferation and of meiotic entry.

### FBFs affect the rates of both stem cell proliferation and differentiation

Here we provide evidence that loss-of-function mutation of *fbf* homologs change the rates of both proliferation and differentiation in *C. elegans* germline SPC. We find that slow proliferation of SPCs in *fbf-2(lf)* is associated with a slower rate of progenitor meiotic entry (differentiation), while the progenitors of *fbf-1(lf)* mutant have a faster rate of meiotic entry (**Figure 8A**). We propose that differentiation and proliferation are simultaneously affected by FBF-mediated control of target mRNAs encoding key molecular regulators of differentiation and cell cycle. Slow meiotic entry rate in *fbf-2(lf)* likely results from translational repression of FBF targets that regulate differentiation; indeed, slower accumulation of FBF target GLD-1 has been documented in this genetic background (Brenner and Schedl 2016). In a similar fashion, mutations of FBF targets *gld-2* and *gld-3* lead to a decrease in meiotic entry rate and to accumulation of excessive numbers of SPCs (Eckmann and others 2004; Fox and Schedl 2015). Conversely, higher meiotic entry rate of *fbf-1(lf)* SPCs might be explained by partial derepression of GLD-1 (Brenner and Schedl 2016; Crittenden and others 2002) and other FBF targets. We find that FBF-2 promotes SPC proliferation through facilitating progression of SPCs through the G2-phase of cell cycle. Thus SPCs of the *fbf-2(lf)* mutant are characterized by longer median G2-phase length. By contrast, the G2-phase of *fbf-1(lf)* SPCs is the same as in the wild type, even though this genetic background shows an increase in the mitotic index. One possible explanation for this observation is that faster meiotic entry rate of *fbf-1(lf)* SPCs depletes the number of progenitors in the pre-meiotic S-phase. Lower total cell number in the distal region then inflates SPC mitotic index. We could not address whether *fbf-1(lf)* germlines have a lower number of progenitors in meiotic S-phase since there are no molecular markers for this developmental stage. Finally, we find that disruption of FBF-mediated regulation of a single B-type cyclin in slowly proliferating and slowly differentiating *fbf-2(lf)* SPCs is sufficient to disturb stem cell homeostasis and leads to excessive SPC accumulation.

### Regulation of Cyclin B by PUF-family proteins in stem cells

PUF mRNA targets have been studied in multiple organisms including *C. elegans*, mouse and human identifying thousands of target mRNAs (Chen and others 2012; Galgano and others 2008; Kershner and Kimble 2010; Morris and others 2008; Porter and others 2019; Prasad and others 2016). One highly conserved group of PUF regulatory targets is related to the control of cell cycle progression. In several developmental contexts stem cells undergo rapid G1/S transitions and spend an extended time in G2, as observed for *C. elegans* germline stem cells and mouse and human embryonic stem cells (Fox and others 2011; Lange and Calegari 2010; Orford and Scadden 2008). PUF proteins facilitate the short G1 phase through repression of proliferation inhibitors such as Cip/Kip family cyclin-dependent kinase inhibitors (Kalchhauser and others 2011; Kedde and others 2010; Lin and others 2019). Additionally, mitotic cyclins B and A are among the core targets of PUF proteins across species including nematode FBFs (Kershner and Kimble 2010; Porter and others 2019; Prasad and others 2016), *Drosophila* Pumilio (Asaoka-Taguchi and others 1999), human and mouse PUM1 and PUM2 (Chen and others 2012; Galgano and others 2008; Hafner and others 2010; Morris and others 2008), and yeast Puf proteins (Gerber and others 2004; Wilinski and others 2015). Cyclin B regulation by PUFs contributes to cell cycle control of *Drosophila* embryonic cell divisions (Asaoka-Taguchi and others 1999; Vardy and Orr-Weaver 2007) and to the control of meiotic resumption during *Xenopus* and zebrafish oocyte maturation (Kotani and others 2013; Nakahata and others 2003; Ota and others 2011). Here, we for the first time report the function of PUF-mediated regulation of mitotic cyclins in the germline stem cells of *C. elegans*. A recent preprint suggests that regulation of cyclin B by PUFs is also observed in mouse embryonic stem cells (Uyhazi and others 2019).

### mRNA deadenylation and PUF-mediated repression

Multiple studies indicate that deadenylation contributes to PUF-mediated translational repression (Goldstrohm and others 2006; Kadyrova and others 2007; Van Etten and others 2012; Weidmann and Goldstrohm 2012). CCR4-NOT deadenylation machinery is conserved in evolution from yeast to humans (Collart and others 2017; Wahle and Winkler 2013). Although deadenylation is required for germline stem cell maintenance in flies, nematodes and mice (Berthet and others 2004; Fu and others 2015; Joly and others 2013; Nakamura and others 2004; Shan and others 2017; Suh and others 2009), the contribution of deadenylation to PUF translational repression *in vivo* is still controversial (Weidmann and others 2014). Here, we find that paralogous PUF proteins FBF-1 and FBF-2 differentially cooperate with CCR4-NOT deadenylation machinery in *C. elegans* germline SPCs.

Multiple lines of evidence suggest that FBF-1’s function *in vivo* is supported by the CCR4-NOT deadenylation. First, the size of germline SPC zone maintained solely by FBF-1 is significantly reduced by a knock-down of CCR4-NOT deadenylase components. Second, FBF-1-mediated repression of FBF target reporter *in vivo* requires CCR4-NOT deadenylase. By contrast, SPC zone maintained solely by FBF-2 and the reporter repression by FBF-2 are not affected by CCR4-NOT component knock down. Taken together, these observations provide genetic evidence that CCR4-NOT promotes FBF-1 function in germline SPCs. The increase in FBF-1 protein levels that we observed after knocking down the CCR4-NOT subunit *ccf-1* (**Figure 3—figure supplement 1C**) might result from the relief of FBF-1 auto-regulation (Lamont and others 2004). Third, both endogenous FBF-1 and GFP::FBF-1 colocalize with a core CCR4-NOT subunit 3xFLAG::CCF-1 *in vivo* by co-immunostaining. An *in vivo* test of protein interaction between GFP::FBF-1 and 3xFLAG::CCF-1 using proximity ligation assay detects positive signal suggesting that these proteins reside in the same complex. By contrast, there’s significantly less *in vivo* colocalization and proximity between GFP::FBF-2 and 3xFLAG::CCF-1. These data are consistent with the idea that FBF-1 and FBF-2 form distinct RNP complexes, of which FBF-1 complex preferentially includes CCR4-NOT deadenylase. Finally, we assessed the length of FBF target poly(A) tail length in the nematodes mutant for each *fbf*, and found that the poly(A) tail length of FBF target *cyb-2.1* was relatively shorter in *fbf-2(lf)* background than in *fbf-1(lf)*. We conclude that FBF-1 selectively cooperates with deadenylation machinery to promote translational repression of target mRNAs (**Figure 8**).

Transcript deadenylation can lead to translational repression or mRNA destabilization (Goldstrohm and Wickens 2008). Measurement of steady-state transcript levels suggested that FBF-1 together with CCR4-NOT decreased the target mRNAs abundance in SPCs. By contrast, FBF-2 promoted accumulation of the target mRNAs. These findings are consistent with the previous qualitative observations that FBF-1 promotes clearance of target mRNAs from the mitotic region of the germline, while FBF-2 can sequester the targets in cytoplasmic foci (Voronina and others 2012). We conclude that in *C. elegans* SPCs mRNA deadenylation primarily results in transcript degradation.

The two FBF proteins are 91% identical in primary sequence (Zhang and others 1997). If FBFs have distinct abilities to engage deadenylation machinery, what are the features of FBF-2 that prevent it from cooperating with CCR4-NOT? PUF RNA-binding domain is sufficient for a direct interaction with the CCF-1 subunit of CCR4-NOT and its homologs in multiple species, including *C. elegans* (Goldstrohm and others 2006; Hook and others 2007; Kadyrova and others 2007; Suh and others 2009; Van Etten and others 2012). However, protein sequences outside of the well-structured RNA-binding domain can promote PUF-induced deadenylation, and are hypothesized to function either through improved recruitment of CCR4-NOT complex or through allosteric activation of CCR4-NOT (Webster and others 2019). We find that the Variable Region (VR) sequences outside of the RNA-binding domain of FBF-1 and FBF-2 play a key role in determining whether these proteins are able to cooperate with CCR4-NOT (Table 2). Mutations of three VRs (VR1, 2, and 4) in FBF-2 result in a protein that now cooperates with CCR4-NOT, suggesting that these regions are necessary to prevent the wild type FBF-2 from engaging with the deadenylase. Conversely, swapping the VR4 of FBF-2 onto FBF-1 renders resulting the chimeric protein FBF-1(vr4sw) insensitive to CCR4-NOT knockdown, indicating that VR4 of FBF-2 is sufficient to prevent cooperation with CCR4-NOT. By contrast, swapping VR3 residing within FBF-2 RNA-binding domain into FBF-1 does not affect the FBF-1(vr3sw) chimera’s cooperation with CCR4-NOT, supporting the importance of protein sequences outside of the RNA-binding domain affecting cooperation with CCR4-NOT. Overall, we conclude that the protein regions outside of the conserved PUF RNA-binding domain regulate the repressive action mediated by each PUF protein homolog. As a result, distinct sequences flanking the RNA-binding domain may lead to differential preference of regulatory mechanisms exerted by individual PUF-family proteins (**Figure 8B and C**). This model provides a foundation for future studies to understand regulatory impact of PUF domain flanking sequences.

**Table 2.** Variable regions outside of RNA-binding domain regulate FBF function

| Transgene | Mutated variable region (VR) sequence | Rescues *fbf-1(lf)*? | Rescues *fbf-2(lf)*? | Dependent on CCR4-NOT |
| --- | --- | --- | --- | --- |
| GFP::FBF-2wt | N/A | No | Yes | No<sup>a</sup> |
| GFP::FBF-1wt | N/A | Yes | No | Yes<sup>a</sup> |
| GFP::FBF-2(vrm) | mutated VR1, 2; VR4 deleted | Yes | No | Yes<sup>b</sup> |
| GFP::FBF-1(vr4sw) | VR4 swapped with FBF-2 | No | Yes | No<sup>b</sup> |
| GFP::FBF-1(vr3sw) | VR3 swapped with FBF-2 | Yes | No | Yes<sup>b</sup> |

## Conclusions

Our results suggest a new model of balancing stem cell self-renewal with differentiation at a population level in *C. elegans* germline. We propose that translational regulation of key mRNA targets by PUF family FBF proteins modulates SPC proliferation together with the rate of meiotic entry or differentiation. Complementary activities of FBF-1 and FBF-2 combine to fine tune SPC proliferation and differentiation to respond to proliferative demands of the tissue. PUF proteins are conserved stem cell regulators in a variety of organisms, and their control of target mRNAs that affect proliferation and differentiation is wide spread as well. The future challenge will be to determine whether PUF-dependent RNA regulation in other stem cell systems might be modulated to adjust stem cell division rate in concert with changing the rate of differentiation.

## MATERIALS AND METHODS

### *C. elegans* culture and strains

All *C. elegans* hermaphrodite strains (supplemental Table S1) used in this study were cultured on NNGM plates seeded with OP50 as per standard protocols (Brenner 1974). All GFP tagged transgenic animals were cultured at 24°C to avoid GFP silencing. Temperature sensitive allele *glp-1(ar202)* is a gain-of-function (gf) mutant and is referred to as *glp-1(gf)* in this study. *glp-1(gf)* is fertile at 15°C, but sterile at 25°C because germ cells fail to enter meiosis and produce tumorous germlines. *glp-1(gf)* was crossed with each single *fbf* loss-of-function (lf) mutant, *fbf-1(ok91)* and *fbf-2(q738),* to generate *fbf-1(lf); glp-1(gf)* and *fbf-2(lf); glp-1(gf).* Double mutant strains and *glp-1(gf)* single mutant were maintained at 15°C. Synchronized L1 larvae of *glp-1(gf)* strains were cultured at 25°C until early adulthood. RNA was extracted from tumorous worms and was subsequently used for qPCR and poly(A) tail length analysis.

### Generation of transgenic animals

All transgene constructs were generated by Gateway cloning (Thermo Fisher Scientific). GFP::FBF-1 and GFP::FBF-2 constructs were generated with the *gld-1* promoter, patcGFP containing introns (Frøkjær-Jensen and others 2016), *fbf-1* or *fbf-2* genomic coding and 3’UTR sequences in pCG150 (Frøkjaer-Jensen and others 2008). GFP::FBF-1(vr4sw) and GFP::FBF-1(vr3sw) constructs were generated similarly with *gld-1* promoter, patcGFP, *fbf-1* genomic coding sequences with swapped variable regions 4 or 3 from *fbf-2*, and *fbf-1* 3’UTR sequences in pCG150. 3xFLAG::CCF-1 construct contains *gld-1* promoter, *ccf-1* genomic coding and 3’ UTR sequences in pCFJ150. 3xFLAG::CYB-2.1wt and 3xFLAG::CYB-2.1fbm constructs contain *gld-1* promoter, *cyb-2.1* genomic coding and 3’ UTR sequences with either wild type (wt, 5’ **UGU**xxxAU 3’) or mutated (fbm, 5’ **ACA**xxxAU 3’) FBF binding sites in pCFJ150.

A single-copy insertion of each GFP-tagged FBF transgene was generated by homologous recombination into universal *Mos1* insertion site on chromosome III after Cas9-induced double-stranded break (Dickinson and others 2013; Wang and others 2016). Similarly, single-copy insertions of 3xFLAG-tagged CCF-1 and CYB-2.1 were generated by targeting universal *Mos1* insertion site on chromosome II. Transgene insertion into universal *Mos1* insertion sites was confirmed by PCR.

### Germline SPC zone measurement

*C. elegans* were synchronized by bleaching, and hatched L1 larvae were plated on NNGM plates with OP50 bacteria or RNAi culture, cultured at specified temperatures and harvested at varying time points depending on the experiment. L1 larvae of *fbf-2(lf); cyb-2.1fbm*, *fbf-2(lf); cyb-2.1wt* and *fbf-2(lf)* were grown at 15°C for 5 days until adult stage. For the time course of SPC zone dynamics, L1 larvae of *fbf-1(lf), fbf-2(lf)* and the wild type (N2) were cultured at 24°C and dissected at 46 hour (late L4 stage), 52 hour (young adulthood) and 63 hour (older adult) time points. In all other SPC zone quantification assays, L1 larvae of all worm strains were cultured at 24°C and dissected for staining at 52 hour time point. Gonads were dissected and stained for mitotic marker REC-8 (Hansen and others 2004), and the length of SPC zone in each germline was measured by counting the number of germ cell rows positive for REC-8 staining before transition zone.

### M phase index measurement

Synchronous cultures of wild type (N2), *fbf-1(lf)* and *fbf-2(lf)* L1 larvae were cultured at 24°C for 52 hours. Gonads were dissected and stained for a mitotic marker REC-8 and an M phase marker phospho-Histone H3 (pH3). Primary and secondary antibodies are described in Supplementary Table S2. M phase index was calculated by dividing the number of pH3-positive SPCs by the number of REC-8-positive SPCs. Percent differences in mitotic indices were calculated through subtracting the mean value of mitotic index of each *fbf* mutant from that of the wild type followed by dividing by the value of the wild type.

### Determination of G2-phase length and meiotic entry rate

G2-phase length analysis and determination of meiotic entry rates were performed by feeding *C. elegans* 5-ethynyl-2’-deoxyuridine (EdU)-containing bacteria as previously described (Crittenden and others 2006; Fox and others 2011; Kocsisova and others 2018), see supplemental materials for details. Germline images were captured as z-stacks spanning the thickness of each germline using a Leica DM5500B microscope. For each replicate time point 7-14 germlines were scored and the data represent 3 or 5 biological replicates. Nuclei were manually counted using the Cell Counter plug-in in Fiji (Schindelin and others 2012) and the Marks-to-Cells R script (Seidel and Kimble 2015) was used to remove multiply-counted nuclei.

Percent differences in median G2-phase length or differentiation rate were calculated as for mitotic index above.

### RNAi treatment

The following RNAi constructs were used: *ccr-4, let-711* (Kamath and Ahringer 2003), *ccf-1* (*cenix:341-c12*; (Sönnichsen and others 2005) and *cyb-2.1* (genomic CDS) in pL4440 (Timmons and Fire 1998). Empty vector pL4440 was used as a control in all RNAi experiments. All RNAi constructs were verified by sequencing. RNAi plates were prepared as previously described (Wang and others 2016) and synchronously hatched L1 larvae were plated directly on RNAi plates, except for *let-711* and *ccf-1(RNAi)*, where L1 larvae were initially grown on OP50 plates and transferred to RNAi plates at the L2/L3 stage. The effect of *cyb-2.1(RNAi)* was confirmed by western blot of 3xFLAG::CYB-2.1. The effectiveness of CCR4-NOT RNAi treatments was assessed by scoring sterility (Figure 3-Figure supplement 1) or embryonic lethality (Supplemental Table 3) in the F1 progeny of the fed animals.

### RNA extraction and RT-qPCR

*glp-1(gf), fbf-1(lf); glp-1(gf)* and *fbf-2(lf); glp-1(gf) C. elegans* were synchronized using bleach, hatched L1s were cultured at 25°C and worms were harvested after 52 hours. Worm pellets were washed 2 times with 1x M9 to remove OP50 bacteria, weighed, flash-frozen using dry ice/ethanol slurry and stored at −80°C. Worm pellets of each strain were collected in triplicate and the qPCR data represent 3 biological replicates. Total RNA was isolated from the worm pellets using Trizol (Invitrogen) and Monarch Total RNA miniprep kit (NEB). RNA concentration was measured using Nanodrop (Thermo Fisher Scientific) or Qubit Fluorometric quantitation (Invitrogen). cDNA was synthesized using the iScript cDNA Synthesis Kit (Bio-Rad) using 1 ug RNA template per each 20 ul cDNA synthesis reaction. Quantitative PCR reactions were performed in triplicate per each input cDNA using iQ SYBR Green Supermix (Bio-Rad) and cDNA diluted 1:10 as template. Primers for *htp-1, htp-2, him-3*, *act-1*, and *tbb-2* were as described (Merritt and Seydoux 2010; Voronina and others 2012). Primers for *cyb-1, cyb-2.1, cyb-2.2*, and *cyb-3* were designed to span exon-exon boundaries to avoid amplification of residual genomic DNA. Abundance of each mRNA in two *fbf* mutants relative to the wild type was calculated using the comparative ΔΔCt method (Pfaffl 2001) with actin *act-1* as a reference gene. After the mRNA abundance of each tested gene was normalized to *act-1*, the fold change values from three replicates were averaged. Finally, fold change values of each tested gene in *glp-1(gf); fbf-1(lf)* and *glp-1(gf); fbf-2(lf)* genetic backgrounds were scaled to the values in *glp-1(gf)* in which the mRNA abundance was set to 1. Differences in mRNA abundance were evaluated by one-way ANOVA statistical tests with linear trend and Tukey’s post-tests.

### <u>P</u>oly(<u>A</u>) tail length (PAT)-PCR

PAT-PCR for the FBF target *cyb-2.1* and control *tbb-2* was performed using a Poly(A) Tail-Length Assay Kit (Thermo Fisher Scientific). RNA templates from *fbf-1(lf); glp-1(gf)* and *fbf-2(lf); glp-1(gf)* strains were the same as used in qPCR analysis. Briefly, G/I tailing, reverse transcription, PCR amplification and detection were performed following the kit protocol. Each G/I tailing reaction used 1 ug total RNA. During PCR amplification, 1 ul of diluted RT sample was used in each PCR reaction and a two-step PCR program was used: 94°C for 2 min, (94°C for 10 sec, 60°C for 1min 30sec) x 35 cycles, 72°C for 5 min. PCR products were assessed using 6% polyacrylamide gel (made with 29:1 Acrylamide/Bis Solution, Bio-Rad) electrophoresis. PCR products were visualized with SYBR Gold stain (Invitrogen) and recorded using ChemiDoc MP Imaging System (Bio-Rad). Poly(A) tail lengths were compared using densitometry analysis in ImageJ.

### Immunolocalization and image analysis

For all immunostaining experiments, *C. elegans* hermaphrodites were dissected and fixed as previously described (Wang and others 2016). All primary antibody incubations were overnight at 4°C and all secondary antibody incubations were for 1.5 h at room temperature. For colocalization analysis of endogenous FBF-1 and 3xFLAG::CCF-1, dissected gonads of *flag::ccf-1* were stained with anti-FBF-1 (Rabbit) and anti-FLAG primary antibodies (Mouse) (Table S2). For colocalization analysis of GFP::FBFs and 3xFLAG::CCF-1, dissected gonads of *3xflag::ccf-1; gfp::fbf-2* and *3xflag::ccf-1; gfp::fbf-1* were stained with anti-GFP (Rabbit) and anti-FLAG primary antibodies (Mouse) (Table S2). Secondary antibodies were Goat anti-Mouse or Goat anti-Rabbit. Germline images were acquired using Zeiss 880 confocal microscope. Localization of FBF granules relative to CCF-1 granules were analyzed in a single confocal section per germline with 4-6 germ cells in SPC zone by Pearson’s correlation coefficient analysis using the JACoP plugin of ImageJ. For each worm strain, 4-8 independent germline images were analyzed and Pearson’s correlation coefficient values were averaged.

### Proximity ligation assay (PLA)

PLA was performed on dissected *C. elegans* gonads following a modified Duolink® PLA Protocol. Fixation was as previously described (Wang and others 2016). Blocking step included incubation in 1xPBS/0.1% Triton-X-100/0.1% BSA for 2x 15 min at room temperature, in 10% normal goat serum for 1 hr at room temperature and in Duolink blocking buffer for 1 hr at 37°C. Primary anti-GFP and anti-FLAG antibodies were diluted in Duolink diluent (Table S2). After overnight incubation with primary antibodies at 4°C, 1:5 dilutions of PLUS and MINUS Duolink® PLA Probes were added to each slide and incubated at 37°C for 1 hr. Next, slides were incubated at 37°C for ligation (for 30 min) and amplification (for 100 min) steps and finally mounted with Duolink Mounting medium with DAPI. Images were acquired using Zeiss 880 confocal microscope. The ImageJ “Analyze Particles” plugin was used to quantify PLA foci in germline images.

### FBF target reporter regulation assay

Reporter transgene with GFP fused to Histone H2B and the 3’ untranslated region (UTR) of *htp-2* (Merritt and others 2008; Merritt and Seydoux 2010) was crossed into *rrf-1(lf), rrf-1(lf)/hT2; fbf-1(lf)* and *rrf-1(lf); fbf-2(lf)* genetic backgrounds. RNAi targeting *let-711* and *ccf-1* were conducted on these reporter strains as described above. The effectiveness of RNAi treatments was assessed by scoring F1 embryo lethality. RNAi treated worms were dissected and fluorescent germline images were acquired on a Leica DFC300G camera attached to a Leica DM5500B microscope with a standard exposure. Percentage of germlines that exhibited target reporter derepression in the SPC zone was scored for each strain.

### Immunoblotting

Synchronous cultures of *C. elegans* were collected at the adult stage by washing in 1xM9 and centrifugation and worm pellets were lysed by sonication. Proteins from worm lysate were separated using SDS-PAGE gel electrophoresis and transferred to a 0.45 μm PVDF membrane (EMD Millipore) as previously described (Ellenbecker and others 2019). Primary and secondary antibodies are described in Supplementary Table S2. Blots were developed using Luminata Crescendo Western HRP substrate (EMD Millipore) and visualized using ChemiDoc MP Imaging System (Bio-Rad).

## ACKNOWLEDGEMENTS

We thank the members of Voronina laboratory for insightful discussions and Geraldine Seydoux for comments on our manuscript. We are grateful to Ella Baumgarten and Jessica Bailey for help with cloning and crosses. We appreciate Ariz Mohammad for sharing the modified R script (originally from the Kimble lab) and instructions on using R for cell counts. Confocal microscopy was performed in the University of Montana BioSpectroscopy Core Research Laboratory operated with support from NIH awards P20GM103546 and S10OD021806. This work was supported by the NIH grants GM109053 to EV and P20GM103546 (S. Sprang, PI; EV Pilot Project PI), American Heart Association Fellowship 18PRE34070028 to XW, and Montana Academy of Sciences award to XW.

## Supplemental Materials and Methods

### EdU labeling

G2-phase length and differentiation rate of germ cells were measured by feeding *C. elegans* EdU-labeled bacteria for varying amounts of time at 24°C (Crittenden and others 2006; Fox and others 2011; Kocsisova and others 2018). Assays for G2 length and differentiation rate were repeated for 3 or 4 times. EdU bacteria plates were prepared by diluting an overnight culture of thymine deficient MG1693 *E. coli* (The Coli Genetic Stock Center; Yale University) 1/25 in 1% glucose, 1mM MgSO_4_, 5 μg/mL thymine, 6 μM thymidine and 20 μM EdU in M9 minimal media. This culture was grown at 37°C for 24 hours, pelleted by centrifugation, resuspended in 10 mL M9 minimal media and plated on 60-mm NNGM plates. Worm strains were synchronized by bleaching, hatched overnight and L1 larvae were cultured on OP50 plates at 24°C for ∼50 hours to reach young adult stage, when they were exposed to EdU-labeled bacteria. After feeding for specified time, worms were picked off EdU plates, dissected on poly-L-lysine treated slides, frozen on dry ice and fixed in ice-cold 100% methanol for 1 min followed by 2% paraformaldehyde/100 mM K_2_HPO_4_ pH 7.2 for 5 min. Next, slides were blocked in PBS/0.1% BSA/0.1% Tween-20 (PBS-T/BSA) for 30 min at room temperature. Samples were incubated with primary antibodies against either phospho-Histone H3 or REC-8 (Supplemental Table 2). After overnight incubation with primary antibody slides were washed 3×10 min with PBS-T/BSA and incubated with secondary antibody for 1.5 hours at room temperature. Secondary antibodies were either Alexa Fluor 594-conjugated goat anti-mouse IgG (H+L) or Alexa Fluor 594-conjugated goat anti-rabbit IgG (H+L) respectively (Supplemental Table 2). After incubation with secondary antibody slides were washed 4×15 min with PBS-T/BSA. Next the Click-iT reaction was performed according to the manufacturer’s instructions (Molecular Probes) with the exception that 2×30 min Click-iT reactions were performed to increase the signal of the Alexa 488 dye. After incubation with the second Click-iT reaction, slides were washed 4×15 min with PBS-T/BSA. Vectashield with DAPI (Vector Laboratories) was added to each sample before cover-slipping. Immunostained germline images were captured as z-stacks spanning the thickness of each germline using a Leica DM5500B microscope for a total of 7-14 germlines per each replicate time point. Nuclei were manually counted using the Cell Counter plug-in in Fiji (Schindelin and others 2012) and the Marks-to-Cells R script (Seidel and Kimble 2015) was used to remove multiply-counted nuclei.

### G2 length and differentiation rate analysis

To calculate the **median duration of G2-phase** animals were fed EdU and collected at 30-minute intervals from 0 to 3.5 hours. Germ cells were co-labeled with anti-pH3 antibody and the fraction of M-phase nuclei that have also completed G2-phase was determined by dividing the number of pH3 and EdU positive nuclei by the total number of pH3 positive nuclei. The percent pH3 and EdU positive nuclei was plotted on the y-axis against the duration of the EdU label on the x-axis and the data were fit to a sigmoidal varying slope curve using GraphPad Prism software, with top and bottom constrained at 100 and 0 respectively (Figure 1—figure supplement 1B). The t_50_ value of the sigmoidal dose-response model was taken as the median duration of G2-phase, or the time at which 50% of pH3 positive cells are also EdU positive.

The **rate of meiotic entry** was calculated by feeding the worms EdU labeled bacteria for 3, 6 or 10 hours and co-labeling the germ cells with anti-REC-8 antibody. The number of nuclei that had entered meiosis in the time since EdU exposure based on being REC-8 negative and EdU positive were counted for each time point. The number of nuclei that entered meiosis was plotted on the y-axis and the duration of the EdU label was plotted on the x-axis in GraphPad Prism software. Linear regression analysis was used to calculate the slope, which corresponds to the number of cells that have entered meiosis per hour.

**Supplemental Table 1.**
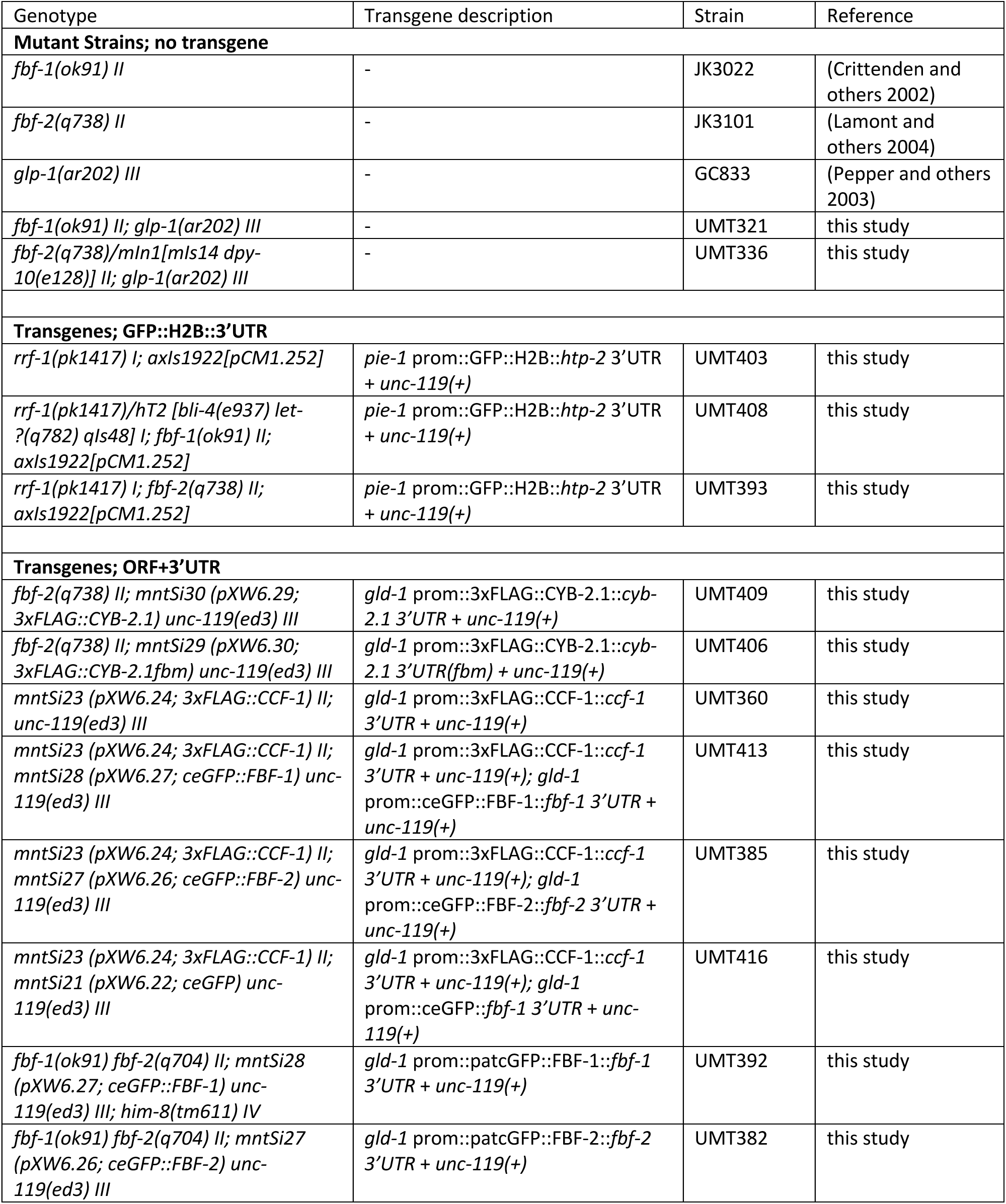
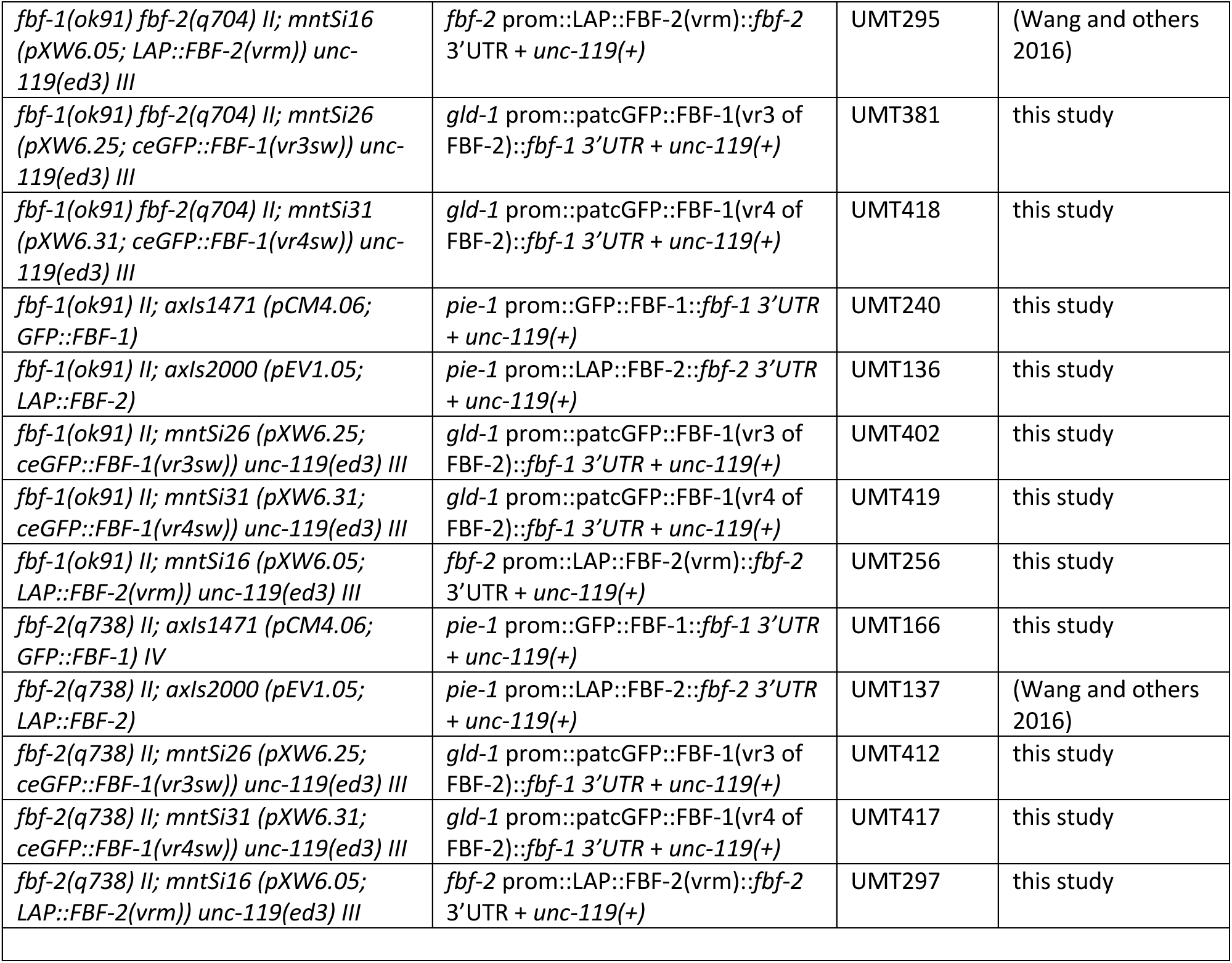
Nematode strains used in this study.

**Supplemental Table 2.** Antibodies used in the study.

| Antibody | Source or Reference | Catalog Number or Designation | Dilution |
| --- | --- | --- | --- |
| Immunostaining, Primary Antibodies |  |  |  |
| Mouse anti-FLAG, IgG1 | Sigma-Aldrich | F1804 | 1:1,000 |
| Rabbit monoclonal anti-GFP, IgG | Thermo-Fisher | G10362 | 1:200 |
| Mouse anti-phospho-Histone H3 pSer10 6G3, IgG1 | Cell Signaling Technology | 9706 | 1:400 |
| Rabbit anti-REC-8, IgG | Novus Biologicals | 29470002 | 1:500 |
| Mouse anti-PGL-1, IgM | DSHB | K76 | 5.2 µg/ml |
| Rabbit anti-FBF-1, IgG | (Voronina and others 2012) | PA2388 | 3.5 µg/ml |
| Immunostaining, Secondary Antibodies |  |  |  |
| Alexa Fluor 594-conjugated goat anti-mouse IgG (H+L) | Jackson ImmunoResearch |  | 1:500 |
| Alexa Fluor 594-conjugated goat anti-rabbit IgG (H+L) | Jackson ImmunoResearch |  | 1:500 |
| Alexa Fluor 488-conjugated goat anti-rabbit IgG | Jackson ImmunoResearch |  | 1:500 |
| Alexa-594 goat anti-mouse IgM | Jackson ImmunoResearch |  | 1:500 |
| PLA, Primary Antibodies |  |  |  |
| Mouse anti-FLAG, IgG1 | Sigma-Aldrich | F1804 | 1:1,000 |
| Rabbit monoclonal anti-GFP, IgG | Thermo-Fisher | G10362 | 1:40,000 |
| Western blotting, Primary Antibodies |  |  |  |
| Mouse anti-FLAG, IgG1 | Sigma-Aldrich | F1804 | 1:1,000 |
| Mouse anti-Tubulin DM1A | Sigma-Aldrich | T6199 | 1:300 |
| Rabbit anti-FBF-1, IgG | (Voronina and others 2012) | PA2388 | 5.2 µg/ml |
| Western blotting, Secondary Antibodies |  |  |  |
| HRP anti-mouse | Jackson ImmunoResearch |  | 1:5000 |
| HRP anti-rabbit | Jackson ImmunoResearch |  | 1:5000 |

**Supplemental Table 3.** Embryo lethality resulting from CCR4-NOT knockdown in the parent generation. The data were obtained in two independent experiments.

| genotype | RNAi | *N* | % Embryo lethality |
| --- | --- | --- | --- |
| wild type | vector | 483 | 0 |
|  | *let-711* | 287 | 49 |
|  | *ccf-1* | 166 | 26 |
|  | *ccr-4* | 556 | 19 |
| *fbf-1(lf)* | vector | 342 | 1 |
|  | *let-711* | 209 | 62 |
|  | *ccf-1* | 127 | 41 |
|  | *ccr-4* | 271 | 20 |
| *fbf-2(lf)* | vector | 361 | 0 |
|  | *let-711* | 375 | 63 |
|  | *ccf-1* | 108 | 77 |
|  | *ccr-4* | 191 | 22 |

**Figure 1—figure supplement 1.**
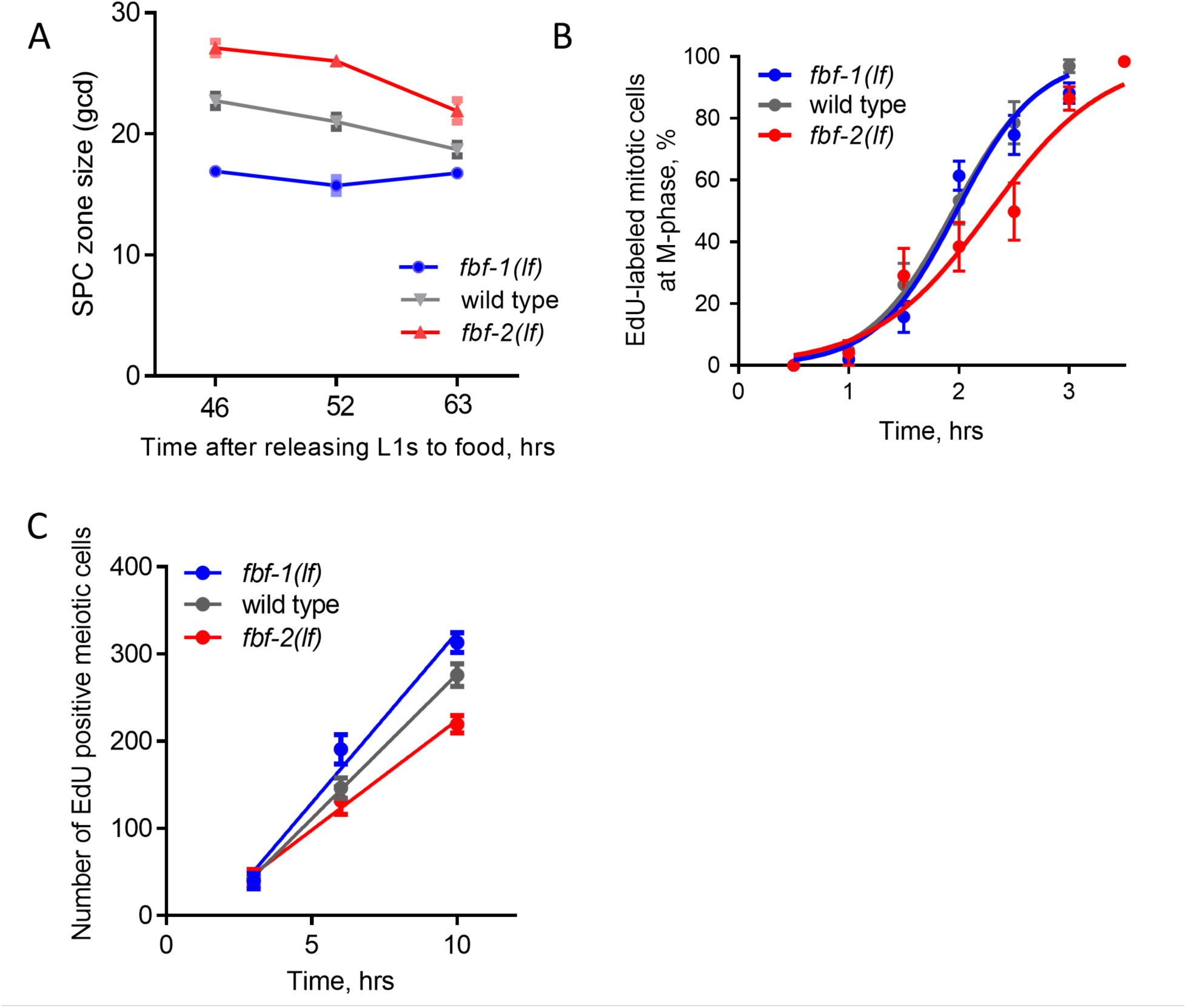
SPC dynamics in different genetic backgrounds. (A) SPC zone length measured as the germ cell diameters (gcd) spanning the stem and progenitor cell zone. X-axis: the time after release of synchronized L1s from starvation at 24_o_C. 46 hrs, L4; 52 hrs, young adult; 63 hrs, older adult. Plotted values are arithmetical means ± S.E.M. 15-20 germlines were scored for each genotype at each time point. (B) Representative time course of EdU labeling of mitotic cells in different genetic backgrounds in one biological replicate. X-axis displays the times when animals were dissected for staining for EdU and pH3. Y-axis indicates the percent of pH3-positive germ cells that are also EdU positive. Plotted values are arithmetical means ± SEM. 9-14 germlines were scored for each genotype at each time point in each individual biological replicate. The median G2 lengths were interpolated as the times when 50% of nuclei in M phase (pH3-positive) were labeled with EdU. (C) Meiotic entry rate of progenitors in different genetic backgrounds. Representative time course of accumulating EdU-labeled, REC-8 negative germ cells in different genetic backgrounds in one biological replicate X-axis displays time points when animals were dissected for staining for EdU and REC-8. Y-axis indicates the number of EdU-positive cells that are negative for REC-8. Plotted values are arithmetical means ± S.E.M. 7-9 germlines were scored for each genotype at each time point in each individual biological replicate. The data were fit to linear regression models, R values were between 0.87 and 0.94. The rates of meiotic entry were determined as the slopes of the regression lines.

**Figure 3—figure supplement 1.**
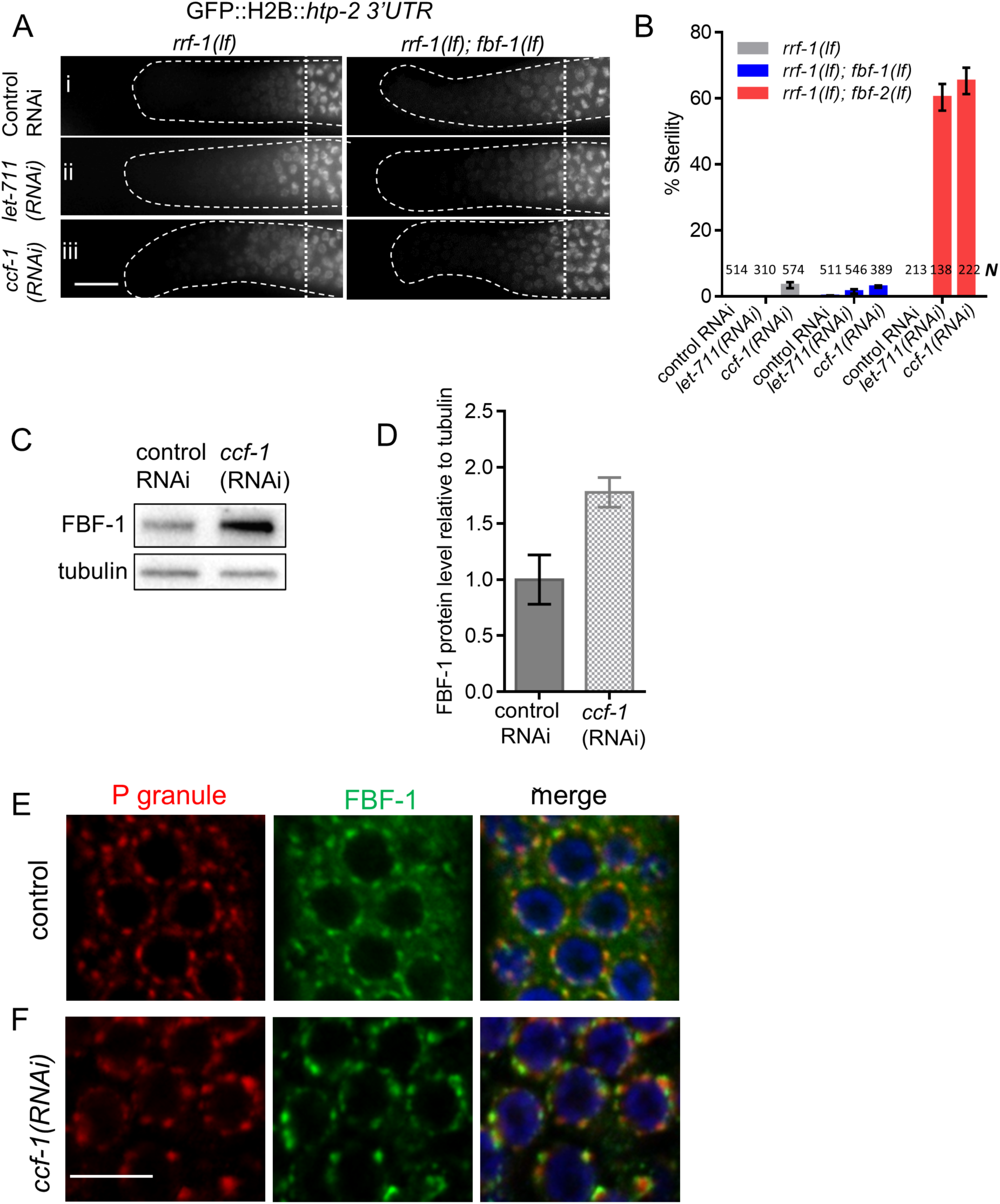
CCR4-NOT deadenylase complex promotes FBF-1 function in germline SPCs. (A) Distal germlines of *rrf-1(lf)* and *rrf-1(lf); fbf-1(lf)* expressing GFP::Histone H2B fusion under the control of the *htp-2* 3’UTR after the indicated RNAi treatments. Genetic backgrounds are noted on top of each image column. Germlines are outlined with dashed lines and vertical dotted lines indicate the beginning of the transition zone. All images were taken with a standard exposure. Scale bar: 10 μm. Efficiencies of RNAi treatments were assessed by sterility (panel B) or embryonic lethality (Supplemental Table 3). (B) Percentage of sterile worms after a knockdown of CCR4-NOT subunits in *rrf-1(lf), rrf-1(lf); fbf-1(lf)* and *rrf-1(lf); fbf-2(lf)* genetic backgrounds. Data were collected from 3 independent experiments. *N*, the number of hermaphrodites scored. (C) A representative Western blot detecting endogenous FBF-1 following *ccf-1(RNAi)*. Tubulin is used as a control. (D) Endogenous FBF-1 protein levels following *ccf-1(RNAi)* determined by densitometry of the western blot results from 3 independent experiments normalized to tubulin. Plotted values are arithmetical means ± S.E.M. (E, F) Confocal images of germline SPC zone co-immunostained for endogenous FBF-1 (green) and P granules (red) in empty vector RNAi control germlines (E) and after *ccf-1* knockdown (F). Scale bar: 5 μm.

**Figure 4—figure supplement 1.**
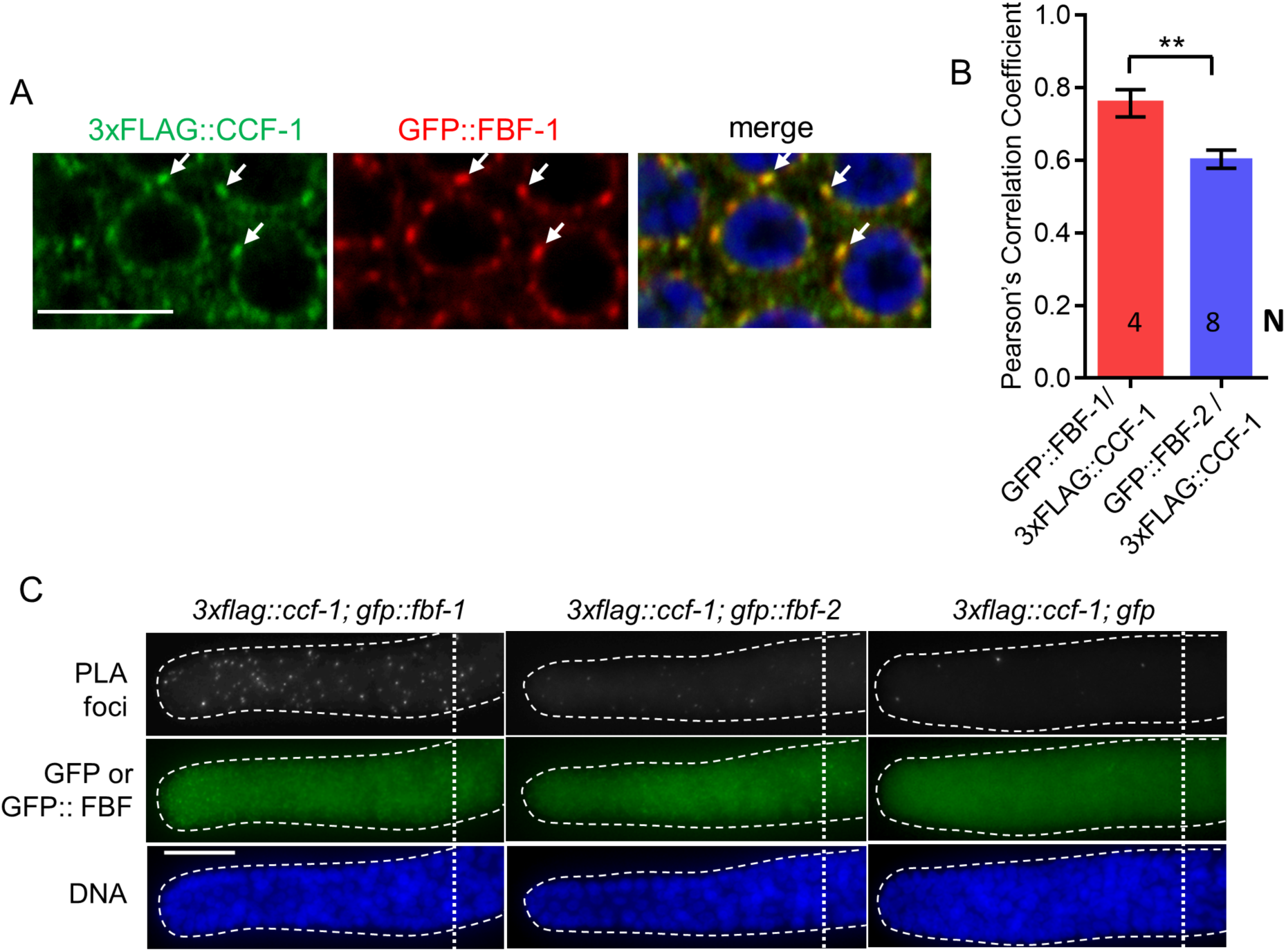
FBF-1 colocalizes with CCR4-NOT complex in germline SPCs. (A) Confocal images of SPCs co-immunostained for GFP::FBF-1(red) and FLAG::CCF-1 (green). DNA staining (DAPI) is in blue. Arrows indicate complete overlap of FBF-1 and CCF-1 granules. Scale bar: 5 μm. (B) Pearson’s correlation analysis of the colocalization between GFP::FBF-1 and FLAG::CCF-1 granules in 4 single confocal sections compared to GFP::FBF-2/FLAG::CCF-1. Plotted values are arithmetical means ± S.E.M. Statistical analysis was performed by Student’s t-test; asterisk marks statistically significant difference, *P*<0.01. The values for GFP::FBF-2/FLAG::CCF-1 colocalization are same as in Figure 4C. (C) Epifluorescent images showing PLA signals (greyscale) and expression of GFP::FBF-1, GFP::FBF-2, and GFP alone (green) in SPCs. DNA staining is in blue (DAPI). Genotypes are indicated above each image group. Scale bar: 10 μm.

**Figure 6—figure supplement 1.**
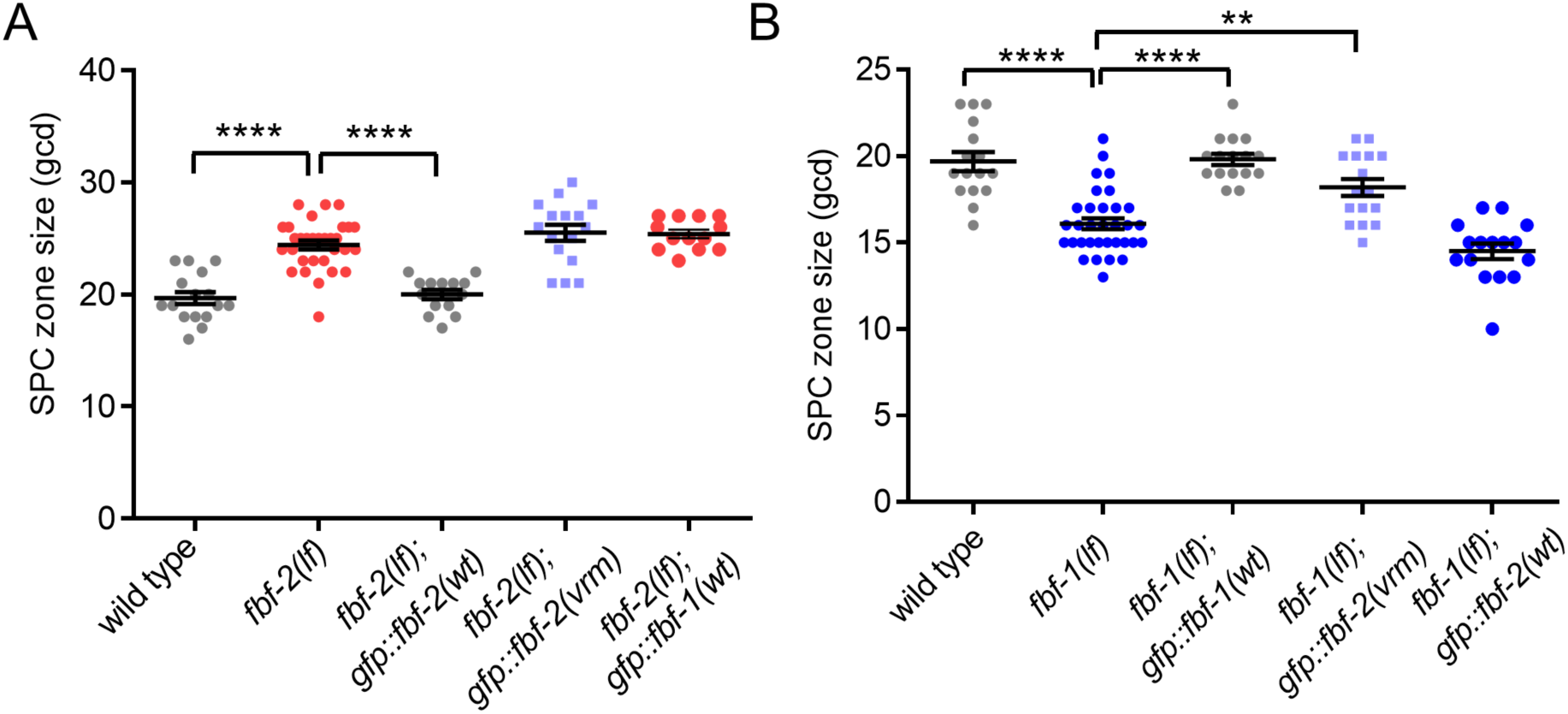
Variable regions 1, 2 and 4 of FBF-2 are required to rescue FBF-2-specific function in germline SPCs. (A) SPC zone sizes were measured after crossing the GFP::FBF-2(vrm), GFP::FBF-2(wt) and GFP::FBF-1(wt) transgenes into *fbf-2(lf)* genetic background. As controls, SPC zone sizes were also measured in *fbf-2(lf)* and the wild type. (B) SPC zone sizes were measured after crossing the GFP::FBF-2(vrm), GFP::FBF-1(wt) and GFP::FBF-2(wt) transgenes into *fbf-1(lf)* genetic background. As controls, SPC zone sizes were also measured in *fbf-1(lf)* and the wild type. (A, B) Plotted values are individual data points and arithmetical means ± S.E.M. Differences in SPC zone size between *fbf-2(lf)* or *fbf-1(lf)* and all other strains in a given group were evaluated by one-way ANOVA test with Dunnett’s post-test; asterisks mark statistically significant differences (****, *P*<0.0001; \*\**P*<0.01). Data were collected from 2 independent experiments and 14-33 germlines were scored for each genotype.

**Figure 7—figure supplement 1.**
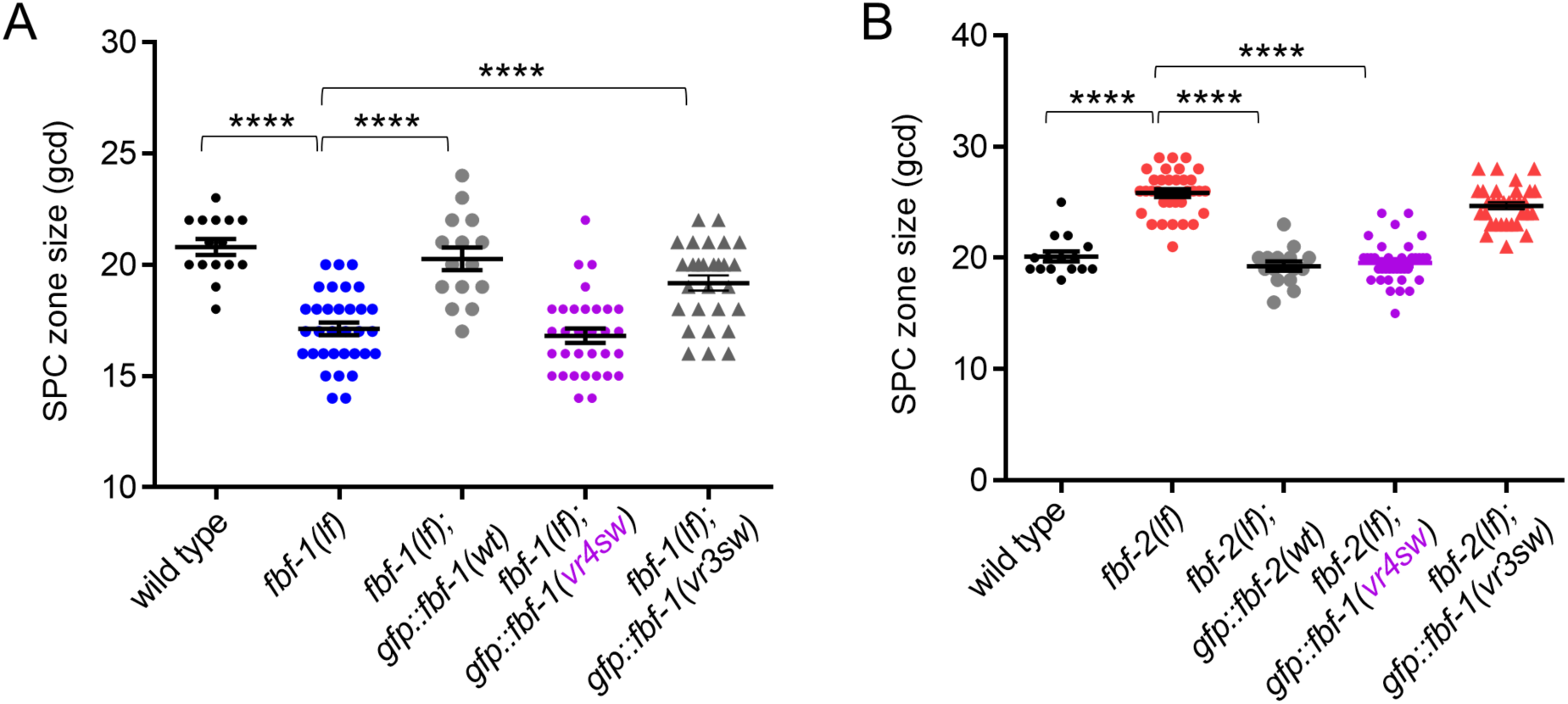
Variable region 4 of FBF-2 allows chimeric FBF-1vr4 to rescue *fbf-2(lf)*. (A) SPC zone sizes were measured after crossing the GFP::FBF-1(vr4sw), GFP::FBF-1(vr3sw) and GFP::FBF-1(wt) transgenes into *fbf-1(lf)* genetic background. As controls, SPC zone sizes were also measured in *fbf-1(lf)* and the wild type. (B) SPC zone sizes were measured after crossing the GFP::FBF-1(vr4sw), GFP::FBF-1(vr3sw) and GFP::FBF-2(wt) transgenes into *fbf-2(lf)* genetic background. As controls, SPC zone sizes were also measured in *fbf-2(lf)* and the wild type. (A, B) Plotted values are individual data points and arithmetical means ± S.E.M. Differences in SPC zone size between *fbf-1(lf)* or *fbf-2(lf)* and all other strains in a given group were evaluated by one-way ANOVA test with Dunnett’s post-test; asterisk marks statistically significant differences (*P*<0.0001). Data were collected from 2 independent experiments and 15-36 germlines were scored for each genotype.

